# Single cell-derived spheroids for real-time growth and metabolomic studies in breast cancer

**DOI:** 10.1101/2025.02.28.638104

**Authors:** Margarida Esteves, Francisca Oliveira, Alexandra Teixeira, Arantxa Ibáñez, Andrea Garcia-Lizarribar, Iratxe Madarieta, José Maria Fernandes, Carolina Rodrigues, Alar Ainla, Lorena Diéguez, Beatriz Olalde, Miguel Xavier, Sara Abalde-Cela

## Abstract

Breast cancer remains a leading cause of cancer-related mortality, with disease progression and metastasis posing significant challenges in treatment. Three-dimensional (3D) cancer models have emerged as valuable tools for studying cancer cell biology in a physiologically relevant microenvironment. Studying the tumour heterogeneity and metabolic adaptations at the single-cell level can be crucial to identify factors driving metastatic progression. Here, we present a novel approach to generate single cell-derived breast cancer spheroids using cell lines (MCF-7 and MCF-10A) within a decellularised adipose tissue extracellular matrix (adECM). Spheroid culture conditions were optimised with integrated plasmonic nanosensors (gold nanostars - GNSs), to enable real-time surface-enhanced Raman scattering (SERS)-based measurements. Our results demonstrated that spheroid growth kinetics and viability in adECM were comparable to commonly used animal-derived matrices, validating its use as a reproducible ECM hydrogel. We further show that the concentration of plasmonic nanosensors used was compatible with cell culture and enabled SERS detection of a model reporter, paving the way for label-free, non-destructive analysis of cancer cell metabolism. This platform offers a promising approach to study cancer progression, including metabolic adaptations, with potential applications in biomarker discovery and preclinical research.

## Introduction

Breast cancer (BC) stands as the second most frequently diagnosed type of cancer at 2.3 million new cases worldwide in 2022 alone. Advances in research, prevention, and healthcare have significantly improved treatment and prognosis. Yet, metastatic BC (mBC) remains incurable, claiming nearly 700,000 lives annually.^1^

Metastasis occurs when cancer cells detach from the primary tumour, enter the blood stream (intravasation), resist anoikis and evade immune surveillance, adhere to capillaries, extravasate, and adapt to the specific microenvironment of a secondary organ. However, only a very small fraction of circulating tumour cells (CTCs) successfully establishes metastatic lesions, which could be explained, at least partly, by the inherent heterogeneity of cancer cells.^2–4^

Tumour heterogeneity manifests at multiple levels – including but not limited to, histological, genetic, or proteomic differences – both within a single tumour (intra-tumoural) and between tumours (inter-tumoural), as well as over time (temporal heterogeneity). Both intra- and inter-tumoral heterogeneity are key players hampering predictions in disease progression and response to treatment.^5–8^

Advances in single-cell analytics, particularly in DNA and RNA sequencing, have revolutionised the way we look at cancer by furthering our understanding of the function and influence of different cell types in carcinogenesis. Indeed, by looking at single cells, one can dissect individual cell parameters that would be otherwise diluted in bulk analyses. Today, single-cell nucleic acid sequencing can be done *in situ*, through the advent of spatial transcriptomics, which enables the resolution of cellular processes while maintaining the spatial context of their specific location within an intact tissue.^9,10^ More frequently, single cell suspensions obtained by being released intact and viable from tissues, are sequenced at high-throughput using microfluidic, droplet-based methods coupled with elegant cellular and molecular barcoding.^10^

Though transforming and incredibly informative, these approaches are still limited in their capacity to study the clonal and temporal evolution of cancer due to restrictions in continuous access to primary tumour samples. In this context, liquid biopsies, for being minimally invasive, offer the opportunity to study cancer evolution through the analysis of tumour biomarkers over time, including circulating nucleic acids, extracellular vesicles, or CTCs. Indeed, in the last decade liquid biopsies have found applicability in cancer screening, treatment selection, assessing treatment responses, and recurrence prediction and detection of minimal residual disease.^11,12^

Yet, cancer studies using single-cell transcriptomics still face two important caveats. One is that cell phenotypes are not always a mirror of their genotype. The other is that the pre-amplification methods still required by some technologies to expand low input nucleic acid contents introduce biases and errors.^13^ While the former highlights the importance of studying cellular parameters other than genomics and transcriptomics, the latter motivates the search for methods that are able to clonally expand the starting cellular material for analysis.^14^

*In vitro* clonal expansion of single cells can be used to mitigate error-prone amplification methods in nucleic acid sequencing, but also contribute to augment the concentrations of metabolites or other cellular materials in multi-omics analyses. *In vitro* clonal expansion can be done in classical 2D cultures or in 3D.^15^ Importantly, unlike cell lines, primary cells (including CTCs) may be difficult to expand in 2D. The success rates of CTC expansion are typically proportional to factors including disease stage, the number of starting CTCs, and the extent of cancer aggressiveness. In addition, their culture often requires media supplemented with growth factors and anoikis inhibitors, hypoxic conditions, co-culture with feeder layers, or low-adhesion culture vessels to prevent adhesion-induced senescence. Despite this, 3D models of human BC have been realised. Cancer spheroids or tumouroids are self-assembled, spherical clusters of cells that mimic the architecture and function of solid tumours more accurately than 2D cultures.^16^ These models are often generated using large numbers of starting cells and through methodologies such as the hanging drop method, the liquid overlay technique, and extracellular matrix (ECM)-based hydrogel cultures, typically involving Matrigel^®^.^17^

The ECM is a critical component of the tumour microenvironment (TME), and in BC, its dynamic and heterogeneous structure have been reported to influence tumour progression and response to therapy.^18^ In cancer research, animal-derived hydrogels containing ECM components like laminin, collagen IV, and proteoglycans are commonly used. However, such matrices are often poorly defined and offer low reproducibility. Alternative ECM-based matrices derived from decellularised tissues have emerged as promising tools for mimicking the biochemical and structural complexity of native ECM in a more reproducible manner.^16,18^

Today, multiomics studies aim to derive associations combining information about cell genotype, phenotype, and even their microenvironment. One of such parameters is cell metabolomics. In cancer, cells reprogram and adjust their metabolic pathways to enhance growth and proliferation, increase their chances of survival, and adapt to a changing microenvironment. Thus, studying characteristic metabolic profiles of cancer cells may be used for earlier cancer diagnosis, selection of patient therapies, or as biomarkers for treatment response.^19^

The standard technique to study CTC metabolomics is mass spectrometry (MS).^20^ In the continuous search for higher sensitivity, other technologies have emerged, which include surface-enhance Raman scattering (SERS) spectroscopy.^21^ SERS stands out as a powerful analytical technique that employs plasmonic metallic nanoparticles to amplify Raman scattering signals, offering the potential of real-time, label-free, and non-destructive analysis with remarkable sensitivity that can arguably reach single molecule detection.^22,23^

In this work, we report a method to obtain single cell-derived BC spheroids while enabling real-time SERS-based analyses. Spheroids were cultured from single BC cells within a decellularized adipose tissue extracellular matrix (adECM), with their growth tracked over time. To ensure both spheroid viability and sufficient signal for SERS measurements, we optimised the concentration of plasmonic nanosensors, here gold nanostars (GNSs), that could be used within the adECM for spheroid growth. Finally, we demonstrated a proof-of-concept for in situ SERS-based metabolite detection within adECM domes.

### Results & Discussion

### 1. Single cell-derived breast cancer spheroids

Tumours exist in dynamic and reciprocal interaction between cancer cells and their TME, which includes non-neoplastic cells, soluble factors, and an altered ECM.^24,25^ Recently, 3D cultures have emerged as promising *in vitro* models to mimic cancer cell interactions with the TME. However, the use of poorly defined animal-derived ECMs hamper control over experimental conditions and reproducibility. Here, we used an ECM from decellularized porcine adipose tissue (adECM). Multiple batches of adECM were obtained with inter-batch variability tested through a series of physicochemical, biochemical, and proteomic analyses (Figure S1a). Remnant DNA was kept at 1.5 ± 0.1 ng/mg, which stands well below the maximum reference values of 50 ng/mg.^26,27^ The protein content was consistent across batches, and cytotoxicity tests confirmed the absence of harmful effects meeting ISO 10993-5 standards. Critically, the proteomic analysis also showed remarkable similarity between the adECM and the original porcine adipose tissue. To assess gelling, we measured the storage (G’) and loss (G’’) moduli of adECM at 10 mg/mL as a function of temperature. At 27°C, the storage modulus surpassed the loss modulus indicating crosslinking and the formation of a solid gel (Figure S1a). The storage moduli of the gelled adECM at 5 mg/mL and 10 mg/mL were 20 and 200 Pa, respectively (Figure S1a), with the latter being on a par with the stiffness of normal breast tissue, and significantly stiffer than the widely used Engelbreth-Holm-Swarm sarcoma-derived matrices at ∼100Pa.^24,26,28^ Envisaging standardised use of the adECM, we studied its stability following sterilisation (with ethylene oxide), storage at −80°C, and prolonged storage at 4 °C (for 3 months). The matrix maintained its stability both as pre-gel and after cross-linking in all conditions tested (Figure S1b), highlighting its robustness for long-term storage and reproducible use in experimental settings.

To initiate spheroid growth, we seeded 40 spatially dispersed BC cells (MCF-7 and MCF-10A) within an 8-µL dome of adECM (Matrigel used as control), with cultures kept for 28 days. A range of ECM concentrations (0.2, 1, 2, 3.5, 5 and 10 mg/mL) were tested to determine the optimal growth conditions. Our data showed no significant differences in both cell proliferation kinetics and viability when the culture media was changed every two or seven days (Figure S2), with a seven-day exchange frequency being thus used for subsequent studies. In these conditions, BC cells grew strictly in adhesion when cultured in ECM concentrations of 1 mg/mL or below (Figure S3a). At intermediate concentrations (2 and 3.5 mg/mL), some cells were able to form spheroids, but the majority still grew in a 2D monolayer (Figure S3b). It was only at 5 and 10 mg/mL that spheroid formation was consistent (Figure 1A). In these conditions, both cell lines tested were able to form spheroids reproducibly at an efficiency rounding 30% of the initially seeded cells (Figure S4). Different media formulations optimised for 3D cultures were not able to increase the efficiency of spheroid formation (Figure S5) and EMEM with 20% FBS supplementation was used in all subsequent studies.

**Figure 1.**
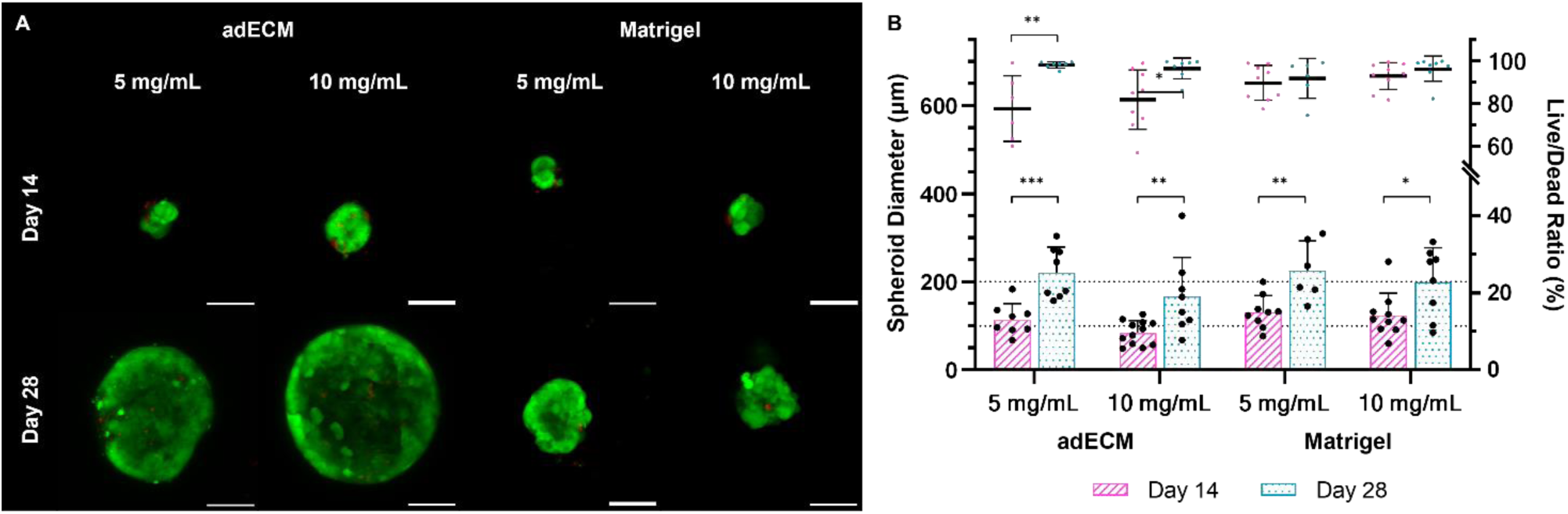
Development of single cell-derived MCF-7 spheroids. A) Live/dead assay (Calcein-AM and BOBO-3) of MCF-7 cells cultured in 8-µL hydrogel domes (Matrigel or adECM at 5 or 10 mg/mL) for 14 or 28 days. Images were acquired using an LSM780 confocal microscope (magnification 20x). B) Spheroid diameters (µm) and live/dead ratio (%) of single cell-derived spheroids cultured in both matrices at concentrations of 5 and 10 mg/mL. Values show mean ± SD (N ≥ 14 spheroids from 3 independent experiments; *p < 0.05; **p < 0.01; **p < 0.001 with p-values obtained using a one-way analysis of variance (ANOVA) with Tukey’s post-hoc test for sample variables that followed a normal distribution (smaller-cell outlet) or a Kruskal–Wallis test followed by a Mann Whitney-U otherwise). (Scale bar: 100 µm)

Critically, Figure 1B shows that the growth kinetics (spheroid diameters at days 14 and 28) and viability (as determined by a Calcein-AM/BOBO-3 Live/Dead assay) of spheroids grown in adECM or Matrigel^®^ were identical. Indeed, at day 28 of culture, spheroids reached a mean size of 187.2 ± 195.2 µm and 204.3 ± 156.7 µm for adECM and Matrigel at 10 mg/mL, respectively, while cell viability remained at 97.0% and 96.9%. These results validate the adECM used in this work as an alternative decellularised 3D matrix that allows tunability to mimic properties of native tumour ECM while minimising batch-to-batch variability. Our current research goals include the adaptation of the current protocols to an adECM of human origin (Figure S1a) aiming to negate interspecies variability, xenogeneic contamination, and ethical concerns related to animal use in research.

### 2. Surface-enhanced Raman scattering spectroscopy of single cell-derived BC spheroids

*In vitro* expansion of single cell-derived tumour spheroids enables clonal interrogation of cancer cell multi-omics and growth kinetics, facilitating studies of inter and intra-tumour heterogeneity that are typically precluded in bulk analyses. Metabolic reprogramming has long been pinpointed as a hallmark of cancer^29^, as first evidenced by Heinrich Warburg, Nobel prize in Physiology in 1931. Thus, it is intuitive that cancer cells that can more effectively reprogram their energy metabolism will become fitter and thus increase their overall chances of survival and consequently their aggressiveness. Adaptations in energy metabolism carry inevitable changes in the cell metabolome. Here, we investigated the potential of SERS spectroscopy for time-resolved tracking of the metabolomics of single cell-derived cancer spheroids.

#### 2.1. Optimisation of growth conditions of BC spheroids in the presence of SERS nanosensors

SERS is a non-destructive, label-free, and ultra-sensitive analytical technique. However, it relies on the use of plasmonic nanostructures for signal amplification. Hence, it was essential to first evaluate the compatibility of spheroid growth in the presence of those plasmonic nanostructures. The nanoparticles of choice were gold nanostars (GNSs), which are sub-100 nm particles with sharp tips (Figure S6) that have shown to provide significant Raman signal enhancement.^30^

Single BC cell-derived spheroids were cultured for 14 and 28 days in the presence of GNSs at 0.5, 2 and 4 mM incorporated in the 10 mg/mL adECM domes on the day of seeding. As a control, PBS (the GNS dispersant) was added in a corresponding volume (Figure S7). A qualitative analysis of cell viability through a live/dead assay showed that there appears to be no effect on cell viability at concentrations of GNSs up to 2 mM (Figure 2A). However, a significant impairment in spheroid growth and overall viability became evident at the highest concentration tested – 4 mM. Despite reaching large sizes by day 28, all spheroids maintained a uniformly spherical shape with a compact core and no necrosis. This is in accordance with studies by Däster *et al*., who reported necrotic core formation only in spheroids exceeding 500 µm – which we did not obtain.^31^ The formation of a necrotic core is also typically associated with a proliferation gradient, with dead or non-proliferative cells in the centre of the spheroid being enclosed by a viable outer layer comprising both quiescent and actively proliferating cells at the periphery.^32,33^

**Figure 2.**
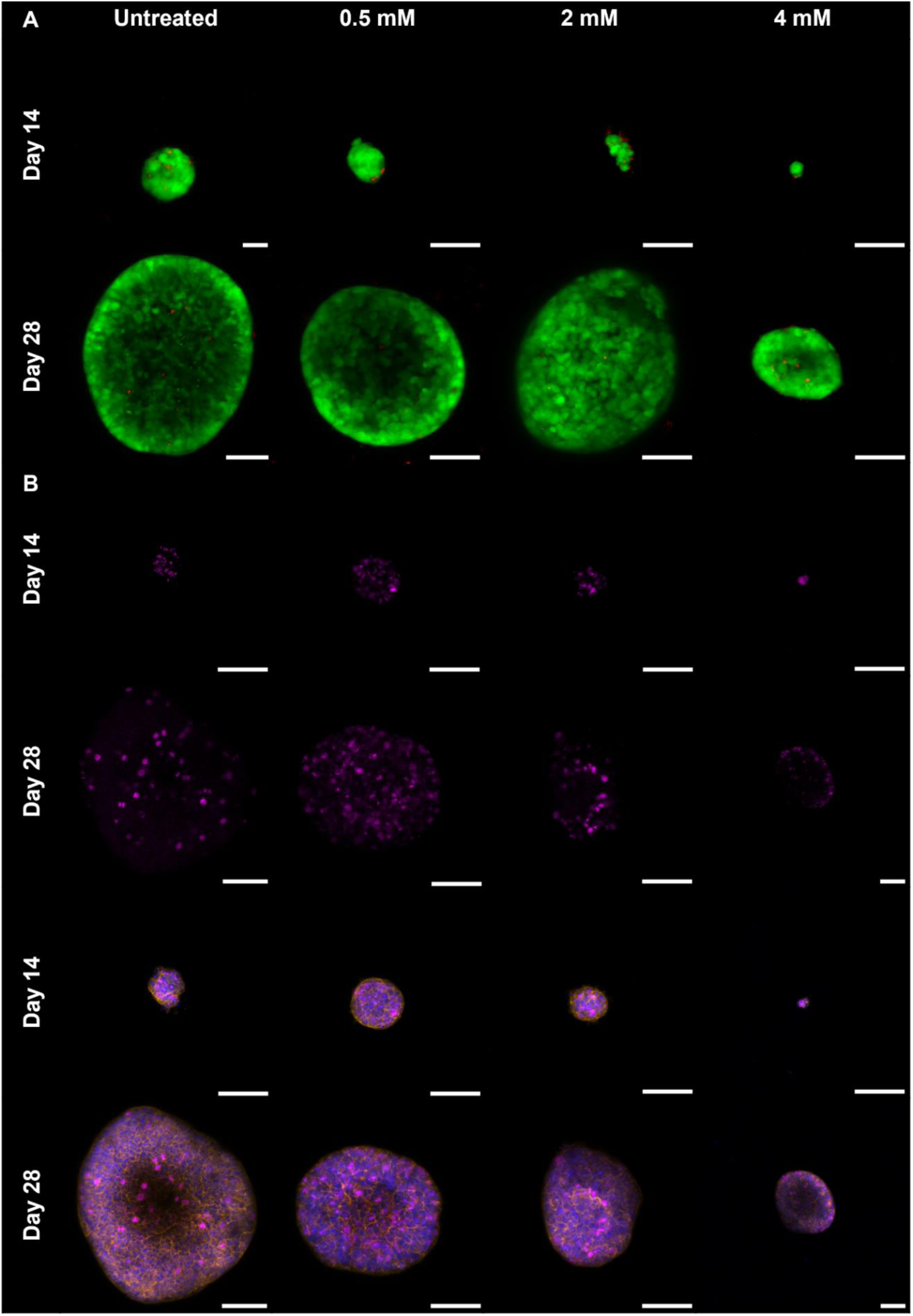
Effect of gold nanostars (GNSs; 0.5, 2 and 4 mM) on the development of MCF-7 spheroids during 28 days of culture. A) Spheroid viability determined by a Live/Dead assay with cells stained using Calcein-AM (live cells, green) and BOBO-3 (dead cells, red). B) Immunocytochemistry of nuclear Ki-67 as a marker of cell proliferation. Cells were counterstained with phalloidin-TRITC and Hoechst 33342 (nuclei – blue, cytoskeleton – orange, and Ki-67 – pink). Images were acquired using an LSM780 confocal microscope (magnification 20x). (Scale bar: 100 µm)

To understand the effect of GNSs on cell proliferation we assessed the expression of Ki-67. Ki-67 is a nuclear DNA-binding protein, which expression varies throughout the cell cycle, reaching its highest levels in G2 and mitosis.^34^ Figure 2B shows a uniform distribution of Ki-67^+^ cells in spheroids at days 14 and 28 at all tested GNS concentrations. Laurent *et al.* reported that spheroids up to 350-µm diameter (as is the case of our spheroids) typically bear a uniform distribution of Ki-67^+^ cells.^35,36^ Spheroids at day 14 appeared to be more proliferative with a higher ratio of Ki-67^+^ cells, which could be anticipated as cells are exposed to less competition over oxygen and nutrients when compared to spheroids at day 28, where cell numbers are significantly increased. The presence of GNSs did not seem to affect Ki-67 expression, which was not in keeping with the growth impairment observed at 4 mM. However, it is important to note that while Ki-67 is a strong indicator of cell proliferation, it is not unequivocal as it has been shown that cells expressing Ki-67 may i) arrest in the G1 phase, ii) proceed to mitosis, or iii) regress to G0.^37^ Thus, we further assessed cell proliferation and spheroid growth using quantitative methods including total DNA quantification and cell metabolic activity by measuring ATP production.

Figure 3 shows quantification of diameters, metabolic activity, and total DNA of MCF-7 spheroids grown for 14 or 28 days in the presence of GNSs. An equivalent analysis was done for MCF-10A cells, which appeared to form smaller spheroids when compared to MCF-7, but these data require further experimental validation (Figure S8). The MCF-7 data evidence the substantial spheroid growth between days 14 and 28, where the mean spheroid diameter of untreated spheroids grew from 89.7 ± 54.1 µm to 342.4 ± 142.8 µm, which was accompanied by a matching increase in the measured metabolic activity and total DNA amount. The total DNA of untreated day-28 spheroids averaged 133.6 ± 67.6 ng, which corresponds to a theoretical number of ∼13,000 cells per spheroid assuming 10.7 pg of DNA per MCF-7 cell (modal chromosome number: 82). This equates to 13 to 14 cell divisions, which is in line with the reported doubling times between 24 and 48h for MCF-7.^38,39^ Importantly, large single cell-derived spheroids of 1,000^+^ cells could enable clonal genomic studies while avoiding whole-genome amplification, which introduces biases and errors including artificial false mutations, allelic dropout, and preferential amplification of one allele.^13^ This is important as it could enable multi-omics studies of the same single cell-derived spheroid – one of our current research goals.

**Figure 3.**
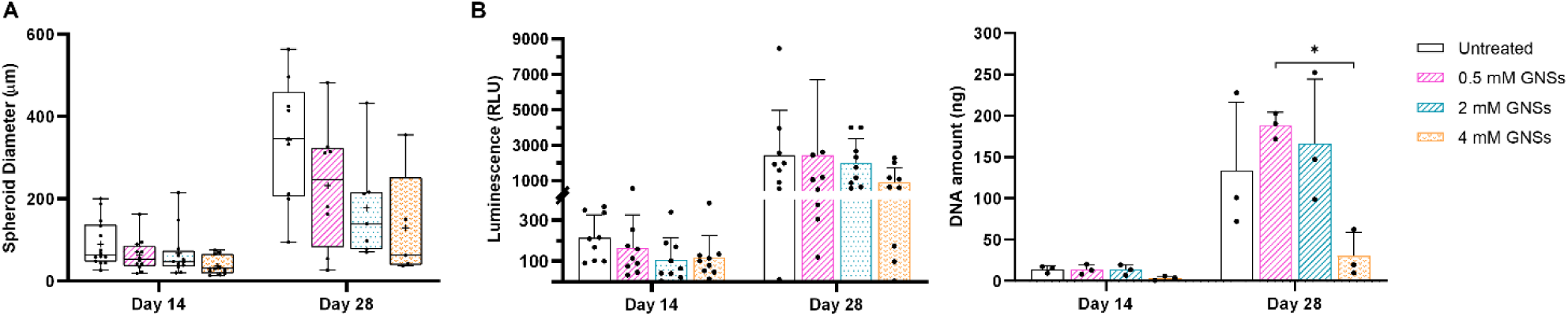
Growth kinetics, metabolic activity and total DNA of MCF-7 spheroids grown in the presence of GNSs (0.5, 2 and 4 mM) at day 14 and 28 of culture. A) Spheroid diameters (µm) of single cell-derived spheroids cultured in the presence of different concentrations of GNSs. Values show mean ± SD (N ≥ 5 spheroids from 3 independent experiments); B) Metabolic activity (RLU) of MCF-7 spheroids cultured in the presence of GNSs. Values show mean ± SD (N = 9 from 3 independent experiments). C) Total DNA amount of spheroids developed adECM with different concentrations of GNSs. Values show mean ± SD (N = 3 from 1 independent experiment; *p < 0.05; with p-values obtained using one-way analysis of variance (ANOVA) with Tukey’s post-hoc test and ANOVA with Kruskal-Wallis multiple comparison tests for normally distributed and non-normally distributed samples, respectively).

This quantitative analysis also validated our observation that GNSs at high concentrations inhibited cell proliferation, which effect was most evident at day 28 of culture. Indeed, when cultured with GNSs at 4mM, spheroids were generally smaller at 128.9 ± 120.9 µm (n.s.), showed slightly lower cell metabolic activity (n.s.) and lower total DNA (p<0.05 *vs* 0.5 mM) when compared to either untreated spheroids or spheroids cultured with lower GNS concentrations. In conclusion, concentrations up to 2 mM of GNSs appeared to be compatible with single cell-derived spheroid growth and were used for all subsequent studies within this work.

### 2.2. SERS metabolomic analysis

Following the determination of the concentration of GNSs that was compatible with spheroid formation and proliferation, we did a proof-of-concept study of *in situ* SERS-based detection of metabolites within the adECM domes. For this, we used a Raman reporter, 1-NAT, following two experimental conditions: i) 1-NAT at 1mM was incorporated inside the domes at the time of plating (Figure 4i); and ii) 1-NAT at 1mM was added to the culture media outside the adECM dome (Figure 4ii). All domes contained GNSs at 2mM. The two conditions were designed to test whether SERS detection would be most efficient in or outside the domes considering the possibility of GNSs leaking out of the domes and the diffusion of cell metabolites to the surrounding culture media.

**Figure 4.**
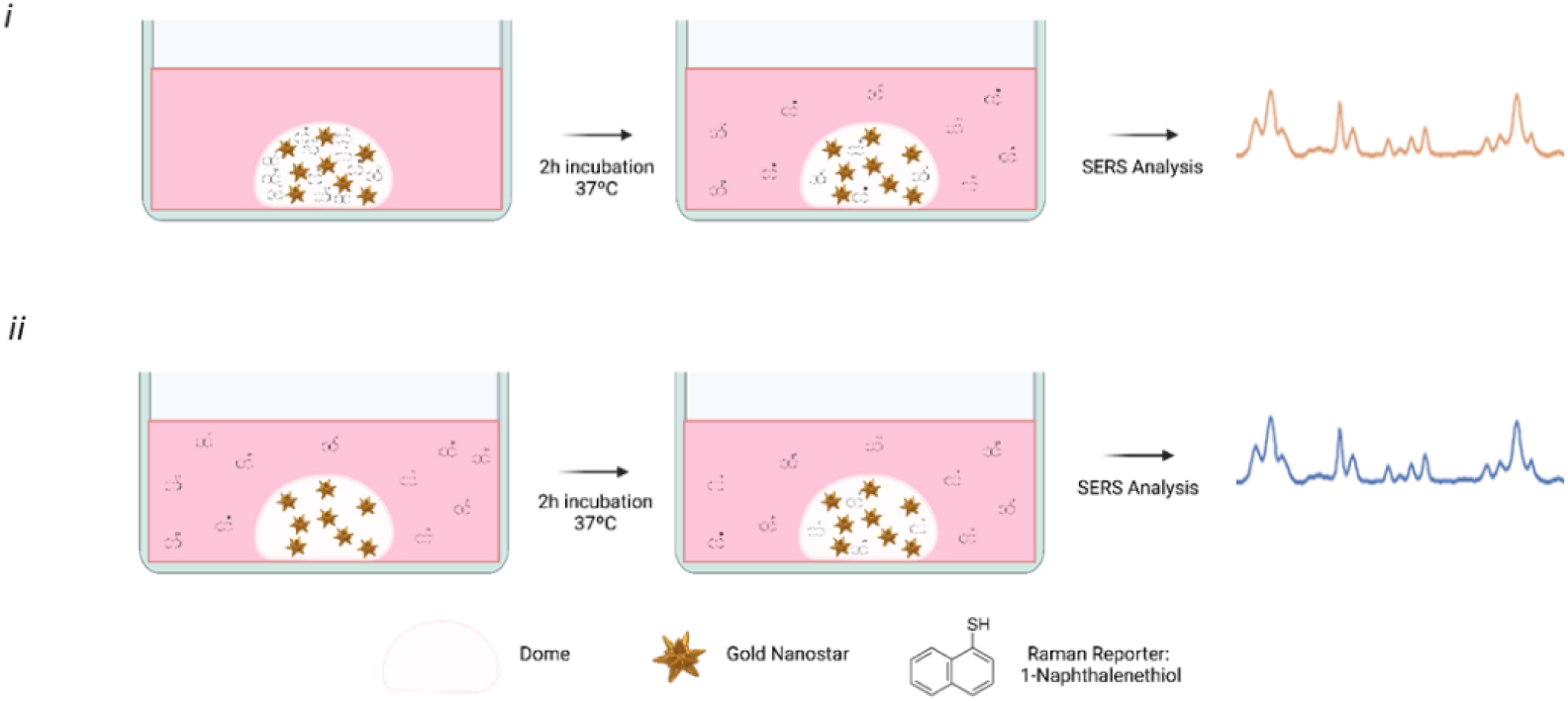
Schematic representation of the experimental process: i) 1-NAT (1 mM) is added within the adECM dome together with GNSs (2 mM), and ii) 1-NAT (1 mM) is added to the media covering the dome (with embedded GNSs. Both conditions allowed diffusion of the Raman reporter, 1-NAT, for 2 hours during incubation at 37°C. SERS measurements were done *in situ* within the dome using an inverted confocal microscope. Note: figure not to scale. Created with BioRender.com

To further assess diffusion through the adECM, we did time-resolved confocal microscopy of fluorescently labelled small (sodium fluorescein; Na-F) and large (70 kDa Dextran-FITC) molecules both from the inside to the outside of the adECM domes, and vice versa. Expectedly, diffusion was faster for fluorescein and dependent on the adECM concentration, with higher concentrations leading to slower diffusion (Figure S9). For example, at 2 mg/mL adECM, the diffusion coefficients of Na-F and Dextran were 1,147 and 200 µm^2^·s^-1^ respectively; and at 10 mg/mL these decreased to 301 and 24 µm^2^·s^-1^. Importantly, even for 70 kDa Dextran and 10 mg/mL adECM, diffusion led to an almost complete concentration equilibrium in and outside of the domes within 60 minutes. This is important as it first shows adequate nutrient/waste product renewal within the adECM matrix throughout the course of culture, and second, shows that cell metabolites will not accumulate within the dome but rather diffuse to the surrounding media over time, which has implications in terms of sensitivity.

It was possible to detect signal from the Raman reporter in both conditions, demonstrating that 1-NAT was able to effectively diffuse from the surrounding culture media to the bulk of the adECM domes where GNSs were present (Figure 5). As a control, we made measurements on positions outside the domes where no signal was detected, indicating that there was no significant leaking of the GNSs from the domes (Figure S10). Both conditions showed a good cross-sectional area of selected 1-NAT characteristic peaks (1,372 cm^−1^), though the intensity varied considerably depending on the position within the dome where the measurement was taken. To show this variation, we did a SERS mapping of regions within the dome (Figure S11). The two mappings highlighted the variable intensity of 1-NAT at 1,372 cm^−1^, but also evidenced the overall higher signal intensity in the condition where 1-NAT was added directly to the adECM dome.

**Figure 5.**
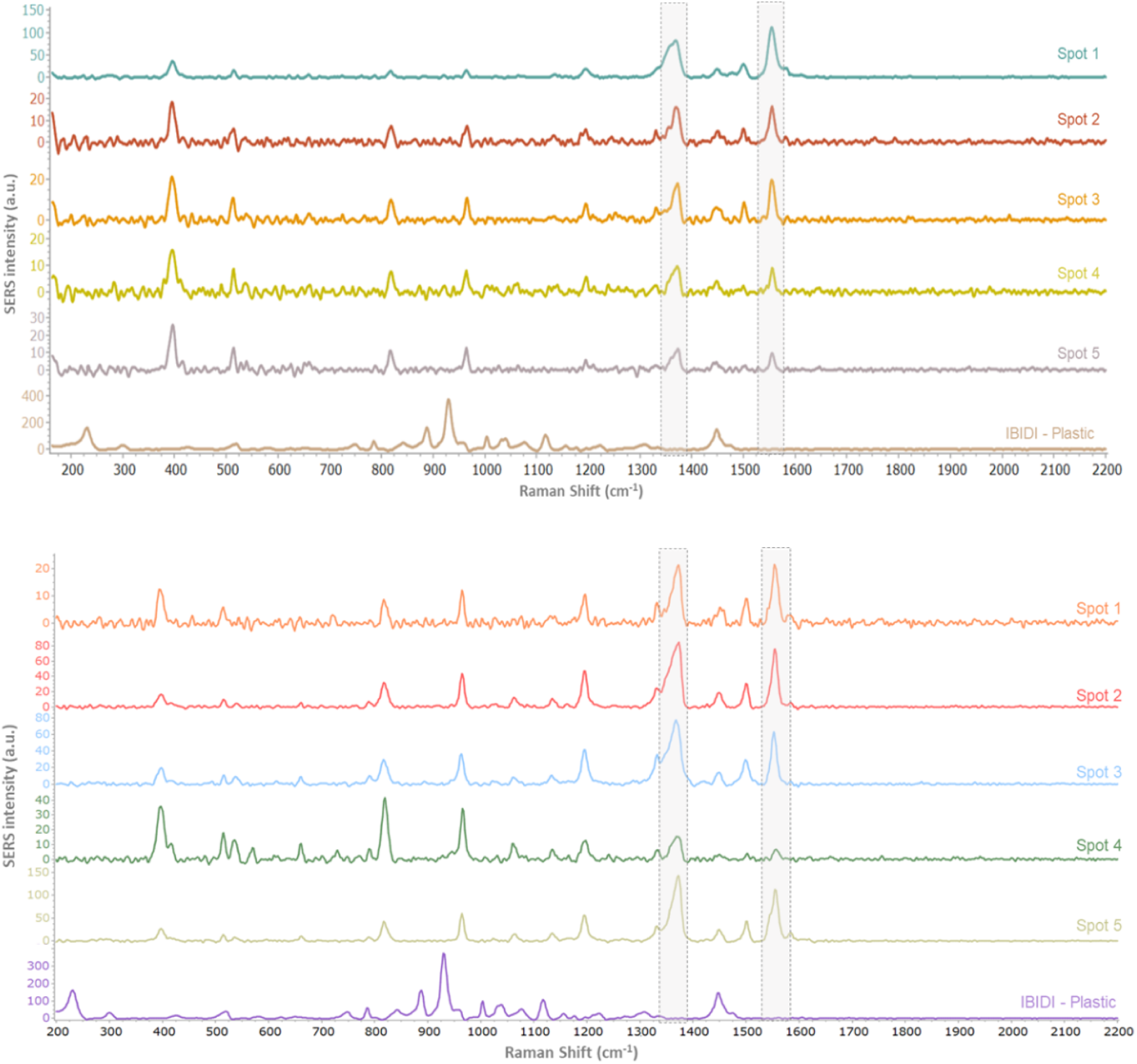
SERS spectra from *in situ* measurements at different position within the hydrogel domes following the experimental conditions i (top) and ii (bottom) respectively. Five different positions were measured and the characteristic peaks of 1-NAT (1,370 and 1,556 cm−1) are highlighted in dashed boxes. All spectra were acquired using an inverted confocal Raman microscope, laser: 785 nm, laser power: 45 mW and with a 50x objective, 2s and 10 acquisitions.

Overall, these data demonstrate the feasibility of our strategy incorporating plasmonic nanosensors within a cell culture hydrogel for time-resolved SERS measurements of cell metabolomics. Building on these early data, our future steps will include SERS characterisation of single cell-derived spheroids looking for potential biomarkers with known intense Raman bands, such as aromatic amino acids like tryptophan, tyrosine, or phenylalanine.^40^ Ultimately, we hope to translate our protocols for label-free, time-resolved metabolomic studies of CTC-derived BC spheroids.

## Conclusion

In this study, we established a method for generating single-cell-derived breast cancer spheroids within a decellularised adipose tissue extracellular matrix (adECM), with the potential to enable real-time SERS-based metabolomic analysis. Our findings demonstrated that adECM supported spheroid formation with growth kinetics and viability comparable to Matrigel^®^. We further showed that the concentration of plasmonic nanosensors used was compatible with cell culture and enabled SERS detection of a model reporter, setting the stage for future *in situ* metabolic profiling. This platform presents a promising tool for studying cancer heterogeneity, metabolic adaptations, and drug responses, with the potential to help bridging the gap between *in vitro* models and clinically relevant insights in cancer research.

## Experimental Methods

### 1. Materials

Dulbecco’s Modified Eagle Medium F12 (DMEM/F12 Cat. number 11320033), Lonza™ Clonetics™ MEBM™ Mammary Epithelial Cell Growth Basal Medium (MEBM Cat. number 11675430), BOBO-3TM Iodide (Cat. number B3586), Calcein-AM (Cat. number B3586), foetal bovine serum (FBS; Cat. number A5256801), bisbenzimide H33342 trihydrochloride (Cat. number 14533), CellTracker Green (Cat. number 10644013), phosphate buffered saline (PBS; Cat. number 10010-023) were acquired from Thermo Fisher Scientific (Waltham, MA, USA). Minimal Essential Medium (EMEM) was acquired from PAN-Biotech GmBH (Aidenbach, Germany). CellTiter-Glo® (Cat. Number G7573) was purchased from Promega (Madison, WI, USA). MammoCult™ Human Medium Kit (Cat. number 05620) was obtained from StemCell Tecnhologies (Vancouver, Canada). Mammary Epithelial Cell Basal Medium (Cat. number PCS-600-030) was obtained from the American Type Culture Collection (ATCC, Manassas, VA, USA). 3D Tumorsphere Medium XF (Cat. number C-28075), Crystal Violet (Cat. number C0775-100G), Fluorescein isothiocyanate–dextran (average molecular weight 4000 KDa Cat. number 46944-100MG-F), Fluorescein isothiocyanate–dextran (average molecular weight 70000 KDa Cat. number 46945-100MGF), fluorescein sodium salt (Cat. number 518-47-8), paraformaldehyde solution 4% (PFA; Cat. number HT5012-1CS), insulin solution human (10mg/mL; Cat. number I9278), TritonTM X-100 (Cat. number T8787) and Trypan Blue solution (Cat. number T8154) were provided by Sigma-Aldrich (St. Louis, MO, USA). Trypsin-EDTA (0.25% trypsin; 0.1% EDTA; Cat. number 15333651) was obtained from Merck Millipore (Burlington, MA, USA). Penicillin/Streptomycin (P/S - Penicillin (10,000 IU/mL) and Streptomycin (10,000 µg/ml); Cat. number P4333) was purchased from Corning (New York, USA). 8-well µ-Slides (Cat. number 80826) were purchased from ibidi® GmbH (Gräfelfing, Germany).

### 2. Cell culture

MCF-7 (HTB-22) and MCF-10A (CRL-10317) were obtained from the American Type Culture Collection (ATCC) and used between passages 6-12 and 5-12 respectively. MCF-7 is a breast cancer cell line representative of the luminal A subtype and MCF-10A is an immortalised mammary epithelial cell line from fibrocystic disease. MCF-7 cells were cultured in EMEM supplemented with 10% foetal bovine serum (FBS), 1% penicillin-streptomycin (P/S), and 0.1% insulin. MCF-10A cells were cultured in DMEM/F12 supplemented with 5% horse serum, 0.5 mg/mL hydrocortisone, 20 ng/mL EGF, 10 µg/mL insulin and 100 ng/mL cholera toxin. The cells were maintained within an incubator set to a humidified environment with 5% CO_2_ at 37 °C. The culture medium was replaced every 2-3 days until the cells reached 70-80% confluency, at which point they were sub-cultured. For subculture, MCF-7 cells were lifted using trypsin-EDTA (0.5 g·mL^-1^ – 0.2 g·mL^-1^) for approximately 10 min and MCF-10A cells using diluted trypsin-EDTA (0.125 g·mL^-1^ – 0.05 g·mL^-1^) for approximately 20 min.

### 3. adECM production

Adipose tissue from porcine and human sources was obtained from a local food company (Jaucha SL, Navarra, Spain) and from Biopredict International, respectively (permission AC-2013-1754, Saint-Grégoire, France), and stored at –20 °C until decellularisation. The decellularisation process, developed and patented by Tecnalia, was modified for human adipose tissue to enhance de-lipidation and improve viscoelastic properties using organic solvents. Tissue preparation involved manual removal of veins and connective tissue, followed by cutting the tissue into 1 cm^3^ pieces. The pieces were washed, blended, homogenized in two stages using different pestles, and centrifuged at 900·g to separate lipids. The supernatant lipids were discarded, and the resulting protein pellets were treated with detergents, solvents, and enzymes. The adECM was then lyophilised and micronised to obtain powder. The powder was finally subjected to enzymatic digestion and pH and salt concentration adjustments to produce *in situ* polymerisable hydrogels.

### 4. 3D spheroid cultures

MCF-7 or MCF-10A cells were harvested and the resulting cell suspension was serially diluted to achieve a final concentration of 40 cells per each 8-µL dome. The cell suspension was then mixed 1:1 with decellularised extracellular matrix from adipose tissue (adECM) at 20 mg/mL. resulting in a final adECM concentration of 10 mg/mL. The suspension was thoroughly mixed and 8-µL domes were pipetted into 8-well ibidi µ-slides. The domes were incubated for 30 minutes at 37 °C to allow adECM polymerisation, after which 250 µL of complete EMEM (with 20% FBS) were added to each well. The cultures were kept in an incubator with humidified atmosphere at 37 °C and 5% CO_2_, with media exchanged every 7 days.

### 5. Cell metabolic activity assay

The culture media was removed and a mixture of 100 µL of CellTiter-Glo^®^ reagent and 100 µL of culture media was added to each well and mixed for 2 minutes on an orbital shaker to induce cell lysis. The plates were then incubated at 37 °C for 10 minutes to stabilise the luminescence signal. Finally, 90 µL from each well were transferred into wells (in duplicate) of a white opaque well plate and luminescence was measured using a BioTek^®^ Synergy H1 microplate reader (Winnoski, VT, USA).

### 6. Live/Dead assay

Cells were rinsed with PBS and then incubated (37 °C; 5% CO_2_; 60 minutes) in a solution containing BOBO-3 Iodide (0.3 µM), Calcein-AM (1.7 µM), and Hoechst 33342 (50 µg/mL) in EMEM supplemented with insulin. The cells were then washed with PBS and maintained in complete EMEM for analysis. As a control for cell death, the cells were incubated with 95% EtOH before staining.

### 7. Immunocytochemistry

Cells were washed with PBS and fixed with 2% paraformaldehyde (pH 7.4) for 20 minutes at 37 °C. The cells were then permeabilised with 0.1% Triton X-100 (v/v) in PBS at RT for 15 minutes and non-specific binding was blocked by incubation with 2% BSA (w/v) in PBS at RT for 60 minutes. Ki-67 was stained by overnight incubation at 4 °C with a rat anti-human Ki-67 antibody (SolA15, eBiosciene™) conjugated with Alexa Fluor™ 488, diluted at 5 μg·mL^-1^ in 0.1% BSA in PBS (w/v). Cells were then washed in 0.1% BSA and counterstained by incubation with Phalloidin-TRITC (0.1 μg·mL^-1^) and Hoechst 33342 (50 μg·mL^-1^) diluted in 0.1% BSA, for 45min at RT.

### 8. Laser scanning confocal microscopy

Images were acquired using a LSM 780 confocal microscope (Zeiss, Oberkochen, Germany) in a controlled atmosphere at 37 °C and 5% CO_2_. Z-stack images were obtained using a 20x lens and a LASOS argon lase (488 nm) and internal diode lasers (405 nm and 561 nm) at a resolution of 1024×1024 pixels. Images were analysed using ImageJ.

### 9. Nucleic acid extraction and quantification

DNA was extracted using the QIAamp^®^ DNA Micro kit following the manufacturer’s recommendations for the isolation of genomic DNA from tissues, and quantified using the Quant-iT™ PicoGreen™ dsDNA Assay Kits and dsDNA reagents also following to the manufacturer’s instructions. Fluorescence (λ_ex_ 480 nm, λ_em_ 520 nm) was measured using a BioTek^®^ Synergy H1 (Winnoski, VT, USA) microplate reader.

### 10. Gold nanostars synthesis and characterisation

Gold nanostars (GNSs) were synthesized using a modified seed-mediated growth method.^41^ Initially, spherical gold nanoparticles (NPs) were produced following the Turkevich method^42^ and coated with polyvinylpyrrolidone (PVP). The PVP-coated gold seeds were collected after centrifugation and stored at RT. For GNS synthesis, PVP was dissolved in dimethylformamide (DMF), and chloroauric acid was added to reduce the Au^3+^ to Au^+^.^41^ Subsequently, PVP-coated gold seeds were added and allowed to react under stirring (90 minutes, 7000 rpm). The resulting GNSs were purified through several washes with isopropanol (IPA) to remove excess PVP and DMF and then stored in IPA protected from light until further use. The synthesized GNSs were characterised by UV–Vis spectrophotometry using a LAMBDA™ 950 UV-Vis-NIR spectrophotometer (PerkinElmer, MA, USA) in a cuvette with a path length of 1 cm. Spectral analysis was conducted over the wavelength range of 300–1200 nm at RT. The GNS morphology was observed by transmission electron microscopy (TEM) using a JEOL2100 (JEOL, Tokyo, Japan) instrument. To prepare the samples, 10 µL of the colloidal GNSs were drop-cast onto a 400-mesh copper grid coated with ultrathin Formvar (TED PELLA, CA, USA) and allowed to air dry. TEM images were captured at 200 kV, and the size distribution analysis was carried out using Image J.

### 11. Estimation of diffusion coefficients

Two experimental conditions (inside-out and outside-in) and two molecules of different molecular weights (70 KDa FITC-Dextran and fluorescein sodium salt) were used to test diffusion within adECM. For inside-out diffusion, adECM was diluted in EMEM containing the test molecules at 100 µM to achieve final adECM concentrations of 2, 3.5, 5, and 10 mg/mL. An 8-μL dome was plated and incubated at 37°C for 30 min, after which 250 μL of culture medium were added allowing diffusion to the surrounding medium. For outside-in diffusion, adECM domes were plated without incorporating the test molecules. Following cross-linking, 250 μL of culture medium containing the test molecules were added to each well to visualise diffusion from the medium into the domes. All observations/acquisitions were done using an LSM 780 confocal microscope (Zeiss, Oberkochen, Germany) acquiring 4×4 tile scan images with a 25s time-lapse interval at 10x magnification.

Diffusion was simulated by finite element modelling (COMSOL Multiphysics 5.4, Sweden) considering the rotational symmetry of the domes and diffusion physics, obtaining a theoretical value for the diffusion coefficient within adECM. Experimental confocal micrographs were saved as series of tiff files and processed automatically using custom scripts written in MATLAB R2018b. Lastly, the experimental fluorescence intensity profile in the center of the dome was fitted to the simulated fluorescence intensity profile, and an experimental diffusion coefficient was obtained using the scaling law D・ t, where D represents the diffusion coefficient and t represents time.

### 12. *In-situ* SERS measurements

SERS measurements were done on crosslinked 8-µL adECM domes containing 2 mM GNSs. In one strategy, 1-NAT at 1mM was mixed directly with the adECM before crosslinking being thus incorporated within the dome. Complete cell culture media was then added to the well plates covering the dome. In another strategy, the adECM domes were seeded without 1-NAT, which was then added also at 1mM to the surrounding culture media. In both cases, the domes and surrounding media were incubated for two hours at 37°C, after which measurements were conducted using an inverted confocal Raman microscope (WITec, alpha300 R*i*) using a 785-nm laser line grating (300 gr·cm^−1^) as the excitation source and a 50× lens. The Raman power was set at 45 mW. SERS spectra were processed with the Project6 WITec software for background, cosmic ray removal and baseline corrections and Spectragryph software for figure preparation.

### 13. Statistical analysis

Results are presented as Mean ± SD. The normality of data was tested using Shapiro-Wilk test with a chosen alpha-level of 0.05. Differences between two group means were compared using the Student’s t-test, while differences between three or more group means were compared using one-way analysis of variance (ANOVA) with Tukey’s post-hoc test and ANOVA with Kruskal-Wallis multiple comparison tests for normally distributed and non-normally distributed samples, respectively.

## Acknowledgements

The authors acknowledge funding from the EU through the 3DSecret project under the HORIZON-EIC-2022-PATHFINDER-OPEN-01-01 programme (grant no. 101099066) and from the UK Research and Innovation (UKRI) council under the UK government Horizon Europe funding guarantee (grant no. 10063360).

## Supplementary data to the manuscript

### S1. Characterization of decellularized ECM from adipose tissue

**Figure S1a.**
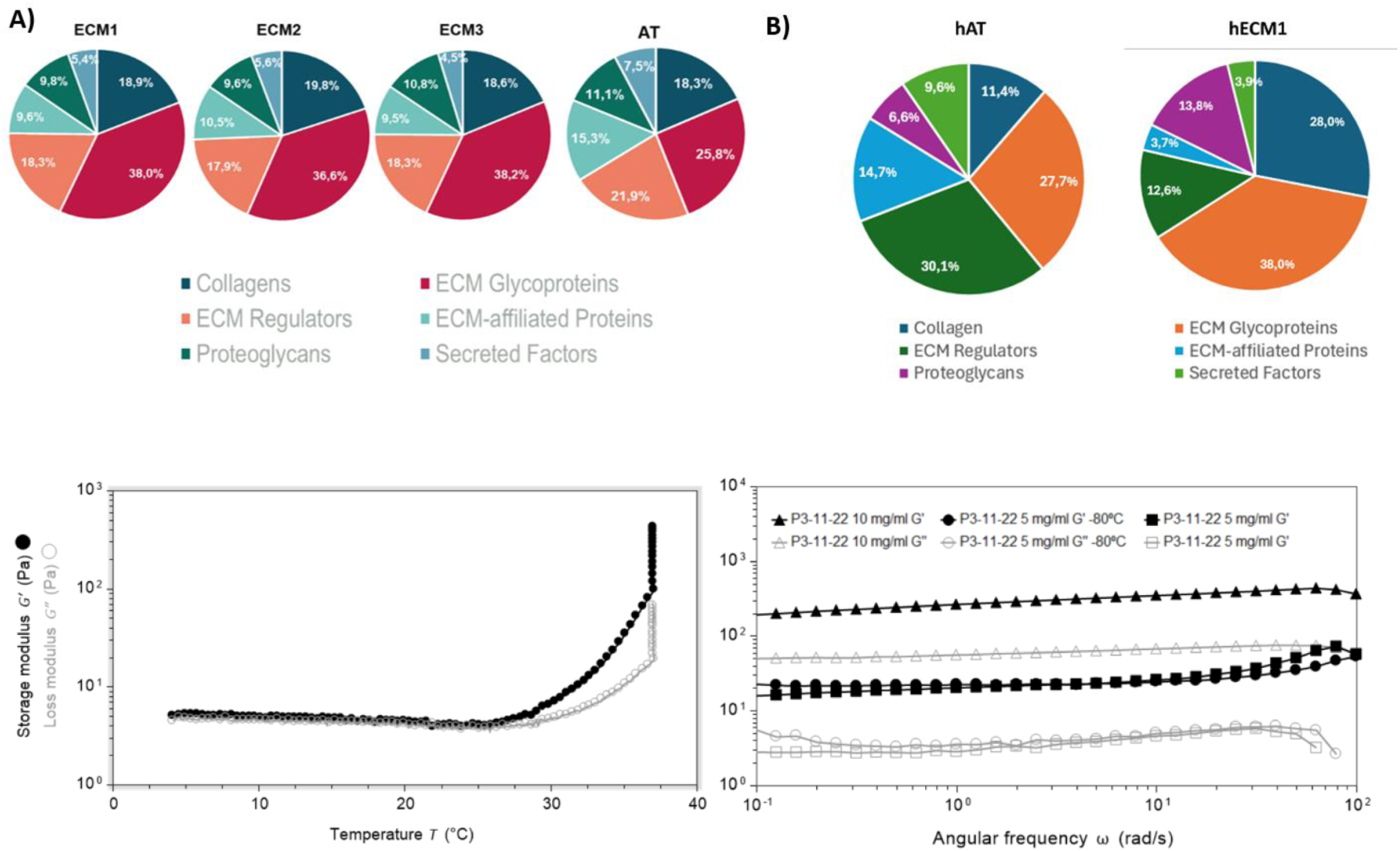
Proteomic analysis of **A)** porcine and **B)** human adECM. **Bottom, left)** Gelling kinetics of 10 mg/ml adECM solutions in a temperature gradient. The storage and loss moduli of adECM at 10 mg/mL were measured as a function of temperature. **Bottom, right) Storage (G’) and loss (G’’) moduli of gelled adECM at 5 and 10 mg/ml concentrations stored at 4 °C or −80 °C.** The effect of material concentration and storage conditions was studied using crosslinked cylinders of adECM tested in a rheometer (TA Instruments H20).

**Figure S1b.**
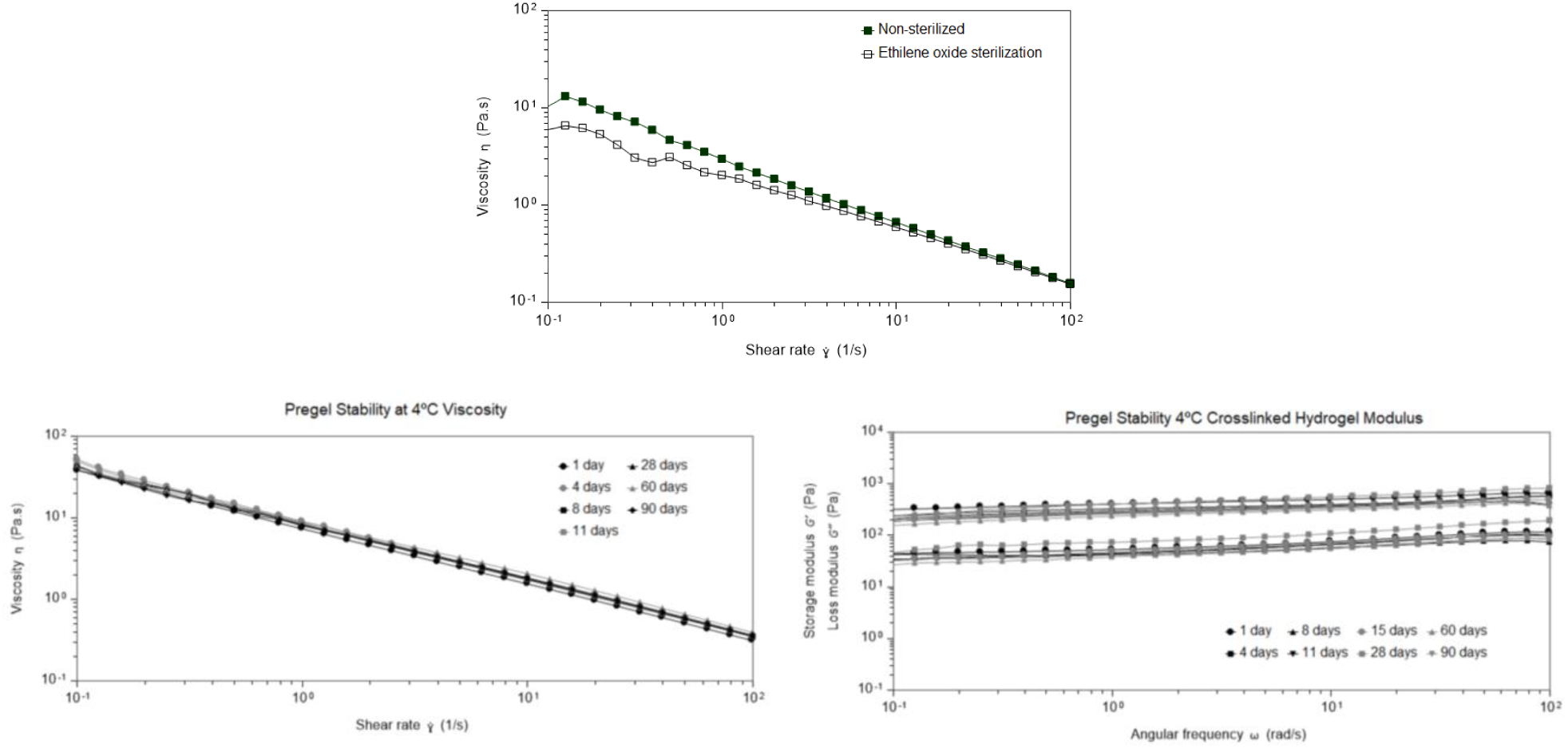
Top) Viscosity measurements at increasing shear rates for sterilized and non-sterilized adECM pre-gels (1.5% w/v). Bottom) Viscoelastic properties at various time points over a total of 90 days of adEC preserved at 4°C: A) Viscosity measurements at increasing shear rates for adECM pre-gel solutions (1.5% w/v). B) Storage (G′) and loss (G′′) moduli of crosslinked hydrogels.

### S2. Media exchange

**Figure S2.**
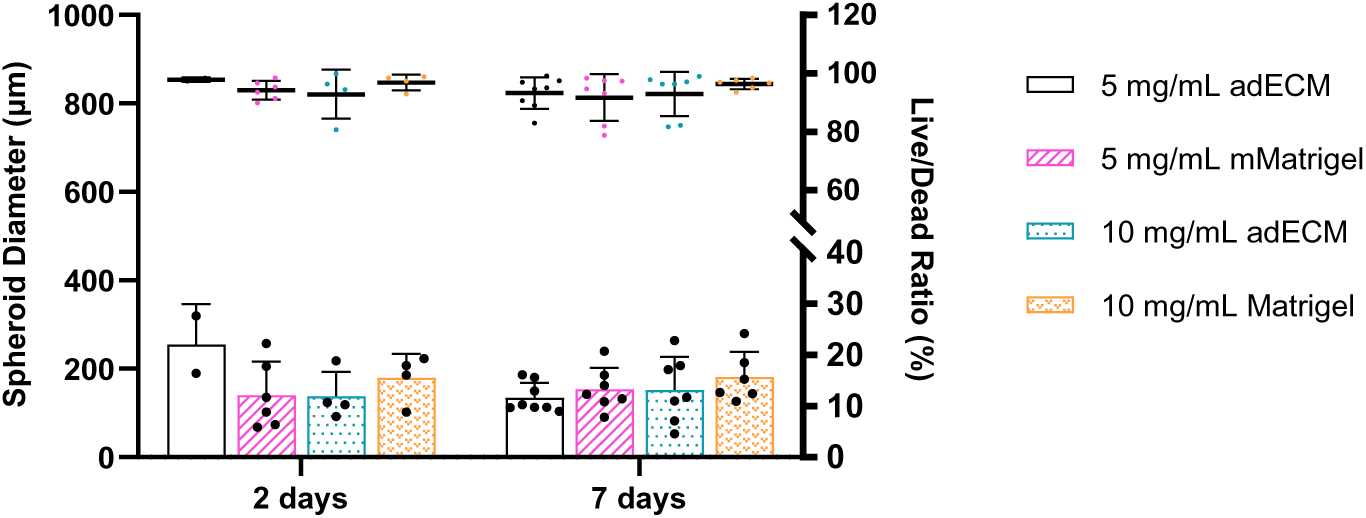
**Effect of media exchange in the development of BC spheroids.** The media was exchanging every 2 or 7 days over 21 days. Spheroid diameters (µm) and live/dead ratio (%) of single cell-derived spheroids.

### S3. Different concentrations of ECM

**Figure S3a.**
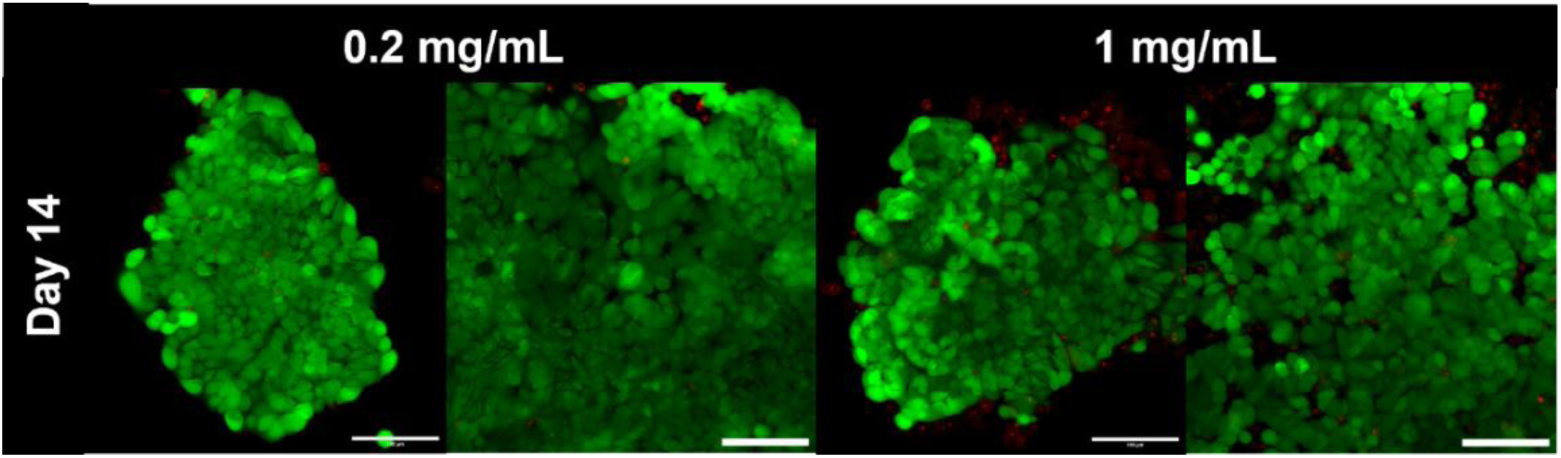
Cells grew adhered at lower concentrations of ECM. Live/dead assay (Calcein-AM and BOBO-3) of MCF-7 cells suspended in Matrigel or adECM at 0.2 or 1 mg/mL and cultured in 8-µL domes for 14 days. Images were acquired using an LSM780 confocal microscope (magnification 20x). (Scale bar: 100 µm)

**Figure S3b.**
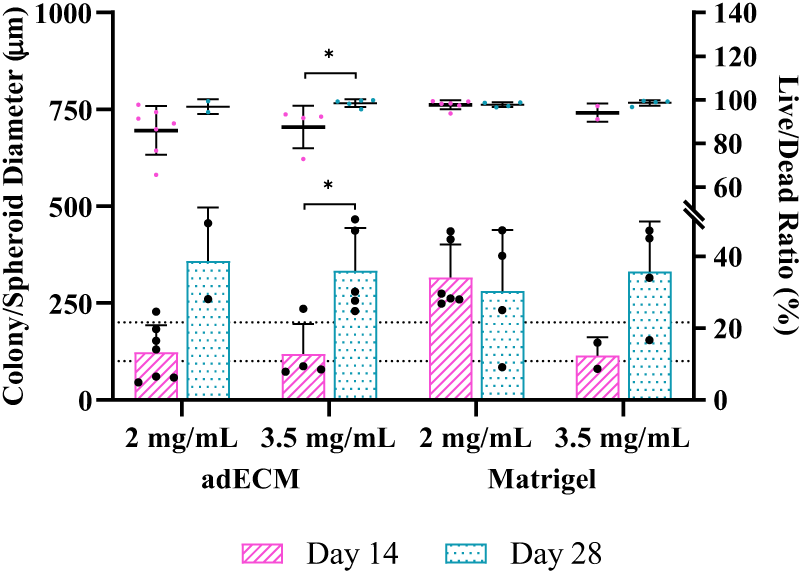
Development of colony/spheroids at intermediate concentrations of ECM. (A) Live/dead assay (Calcein-AM and BOBO-3) of MCF-7 cells suspended in Matrigel or adECM at 2 or 3.5 mg/mL and cultured in 8-µL domes for 28 days. Images were acquired using an LSM780 confocal microscope (magnification 20x). b) ). (B) Colony/Spheroid diameters (µm) and live/dead ratio (%) of single cell-derived spheroids cultured in both matrices at concentrations of 2 and 3.5 mg/mL.

### S4. Seeding efficiency and % spheroid growth

**Figure S4.**
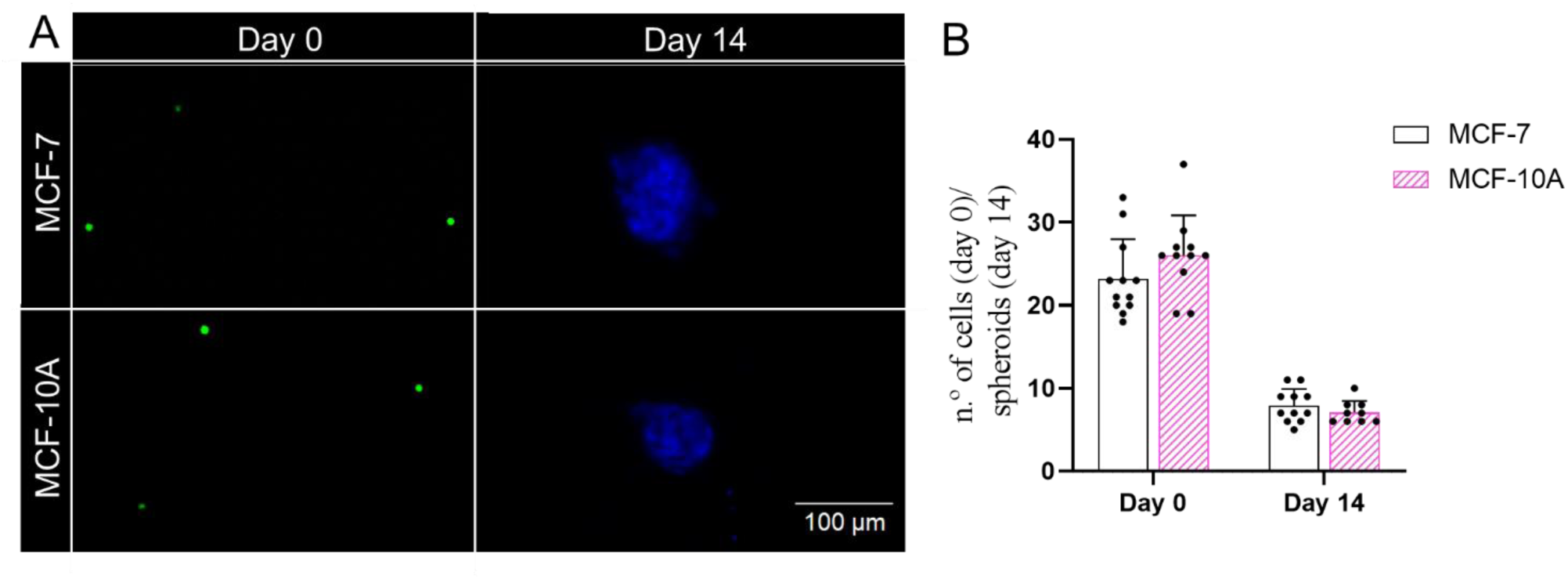
**Number of MCF10-A and MCF-7 cells seeded inside each adECM and number single-cell derived spheroids at day 14.** (A)The number of CellTracker Green stained cells seeded inside each dome was counted on day 0 and the number of Hoechst-stained spheroids inside each dome was counted on day 14 using a LSM780 confocal microscope (magnification 20x). (B) shows the data quantification as mean ± standard deviation. nsp > 0.05 were calculated using a Student’s t-test at day 0 and day 14 (n= 4 replicates from three independent experiments).

### S5. Effect of different culture media in the development of single cell-derived cancer spheroids

**Figure S5.**
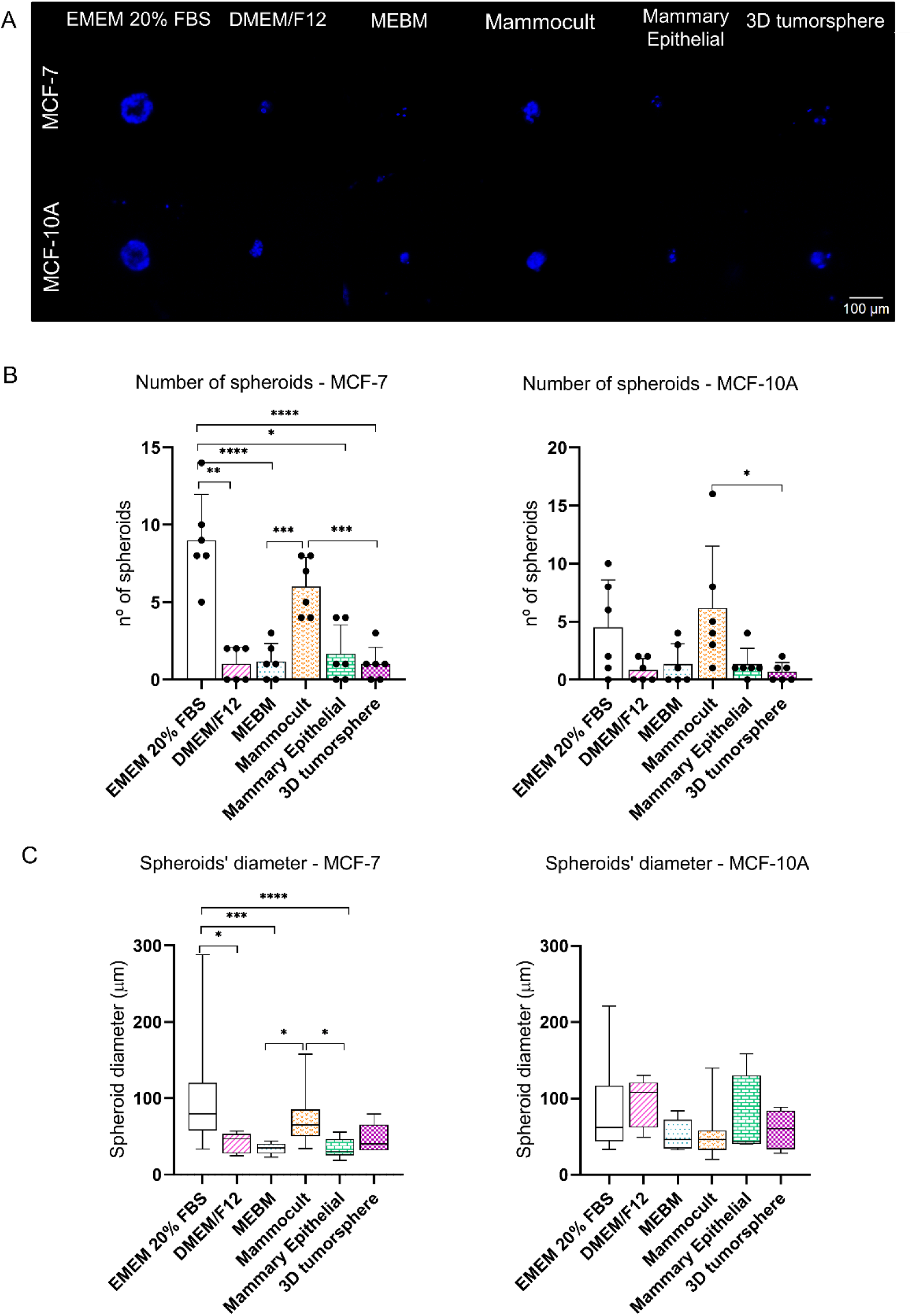
3D colony formation assay of MCF-7 and MCF-10A cells in different culture media. (A) Spheroids were stained with Hoechst and the images were acquired using an LSM780 confocal microscope (magnification 20x). The number (B) and diameter (C) of the spheroids were assessed using ImageJ. Data is shown as mean ± standard deviation. *p < 0.05; **p < 0.01; ***p<0.001; ****p < 0.0001 were calculated using a mixed effects analysis due to non-normally distributed samples. (n=2 replicates from three independent experiments).

### S6. Gold nanostar characterisation

**Figure S6.**
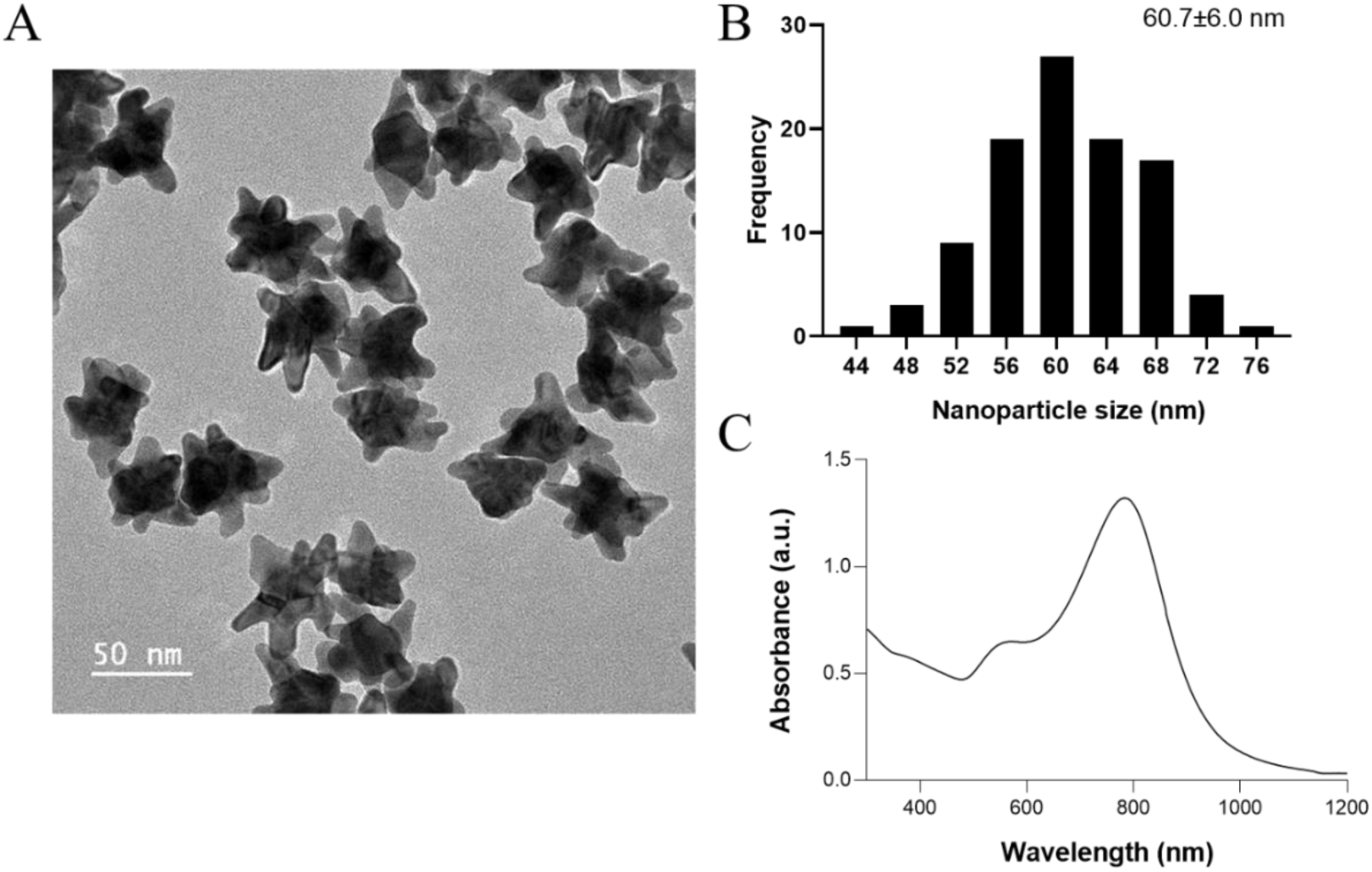
Morphological and optical characterisations of the GNSs. (A) TEM image of the GNSs; (B) Histogram representing the size distribution of the obtained GNSs. For this, 100 GNSs were measured using ImageJ and the average size of the GNSs was 60.7 nm with a standard deviation of 6.0 (60.7±6.0) nm; (C) UV-vis spectrum of GNSs. The plasmon band located at 782 nm is related to the tips of the GNSs. The peak located at a maximum wavelength of 571 nm appears as a result of the core of the GNSs.

### S7. PBS and Negative Control of MCF-7 cells in viability assay

**Figure S7.**
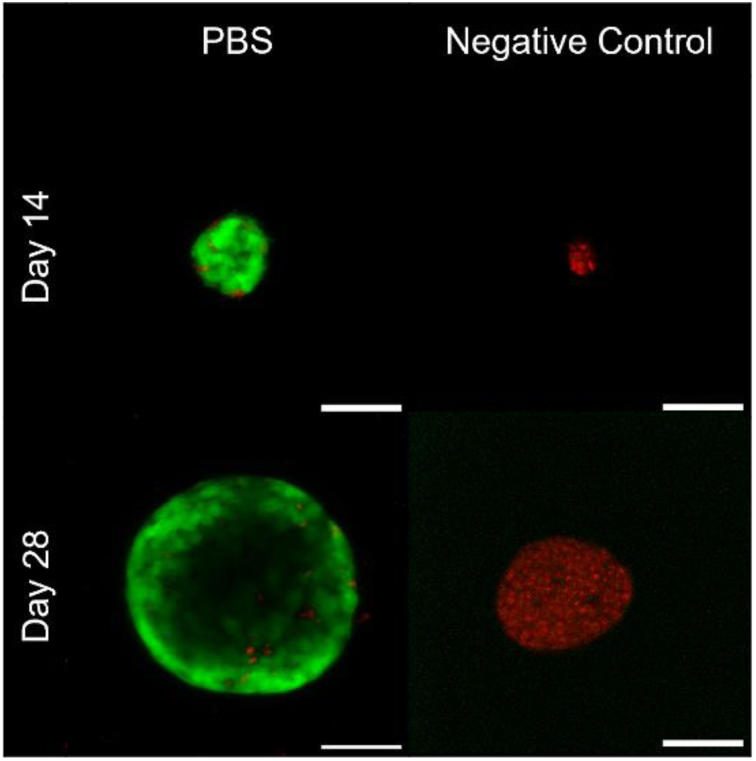
Spheroid viability at day 14 and 28 of culture to PBS (used as a GNSs dispersant – control) and negative control of the Live/Dead assay. Images were acquired using an LSM780 confocal microscope (magnification 20x). (Scale bar: 100 µm)

### S8. MCF-10A

**Figure S8.**
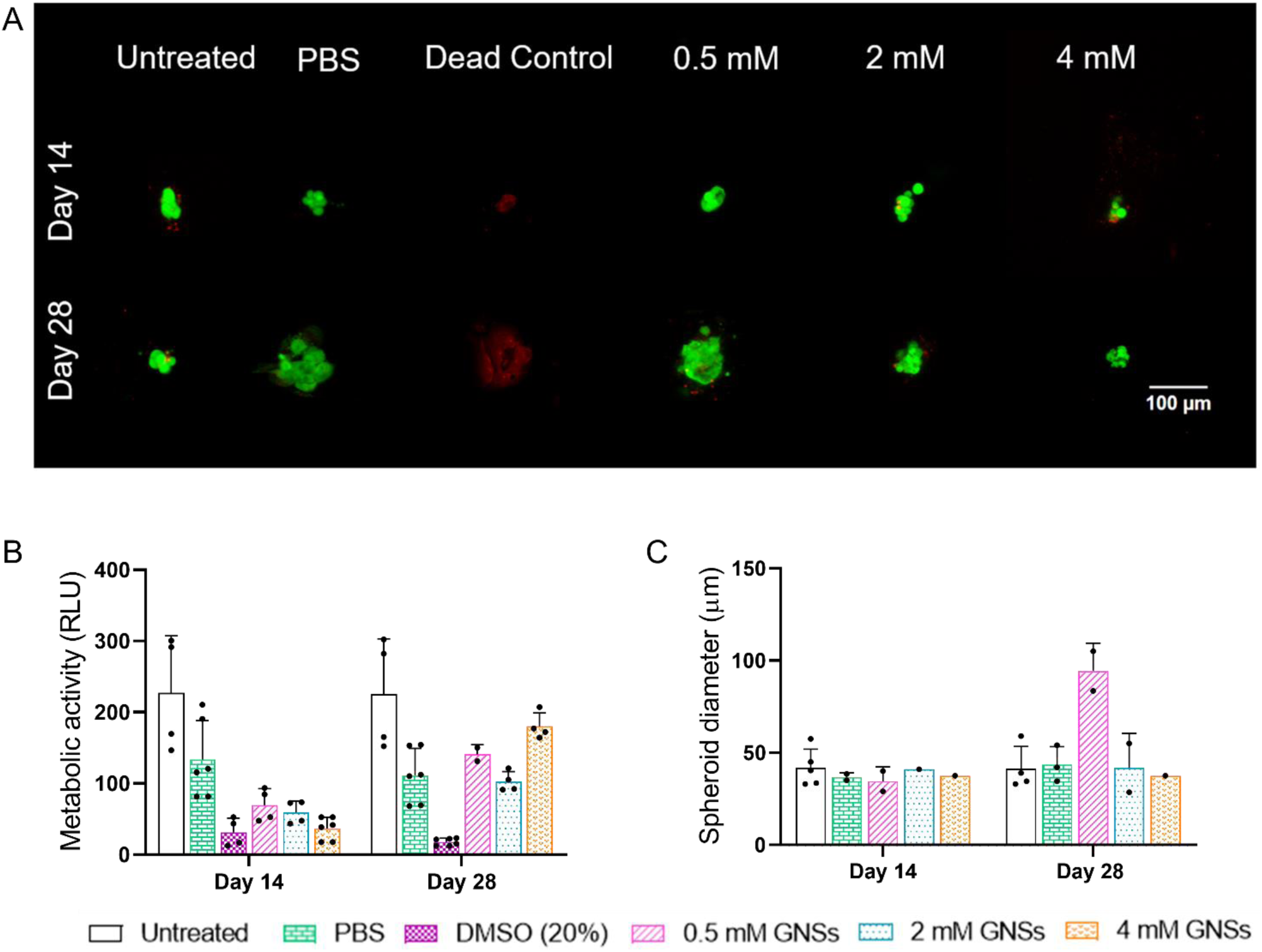
(A) Cell viability of MCF-10A spheroids in adECM domes with different concentrations of GNSs (0.5, 2 and 4 mM) at days 14 and 28. Cells were stained with Hoechst (nuclei), calcein-AM (live cells) and BOBO-3 iodide (dead cells), and the images were acquired using an LSM780 confocal microscope (magnification 20x). (B) Metabolic activity of MCF-10A spheroids in adECM domes with different concentrations of GNSs (0.5, 2 and 4 mM) at day 14 and 28 measured by the CellTiter-Glo assay. Data is shown as mean ± standard deviation. (C) Spheroid diameter of MCF-10A spheroids in adECM domes with different concentrations of GNSs (0.5, 2 and 4 mM) at day 14 and 28 measured using the ImageJ software.

### S9. Diffusion

**Figure S9.**
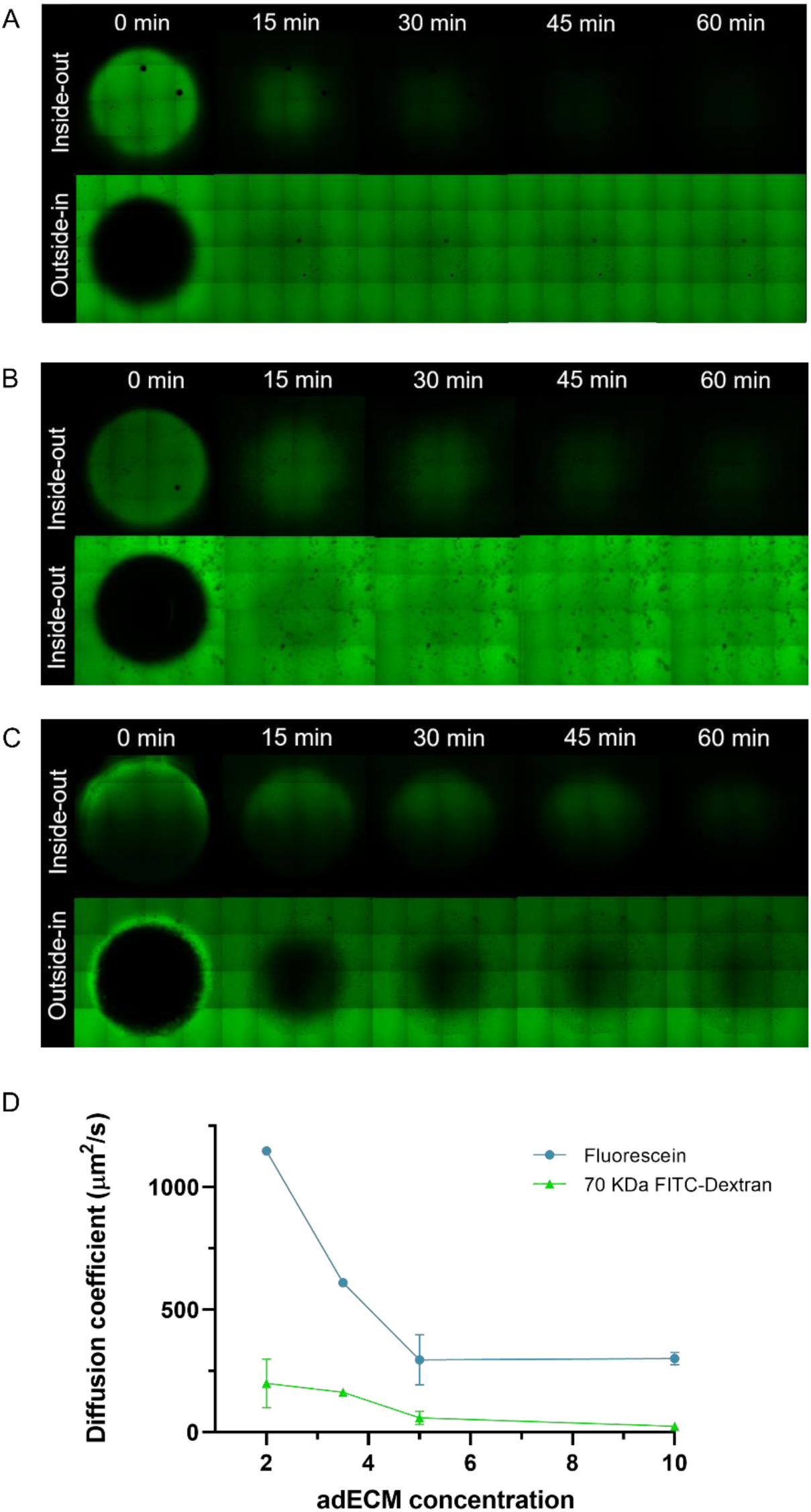
Complete diffusion of fluorescein (A), 4 KDa Dextran (B) and 70 KDa Dextran (C) molecules through 10 mg/mL adECM domes. Quantification of diffusion coefficient of different sized molecules through adECM domes (D). Experimental data was processed using a MATLAB script to determine the fluorescence intensity in the center of the dome through time. Then, experimental data was fitted to a simulated model of diffusion through the adECM domes performed using COMSOL Multiphysics 5.4 software, calculating the diffusion coefficient. Images were acquired using an LSM780 confocal microscope (magnification 10x).

### S10. SERS controls

**Figure S10.**
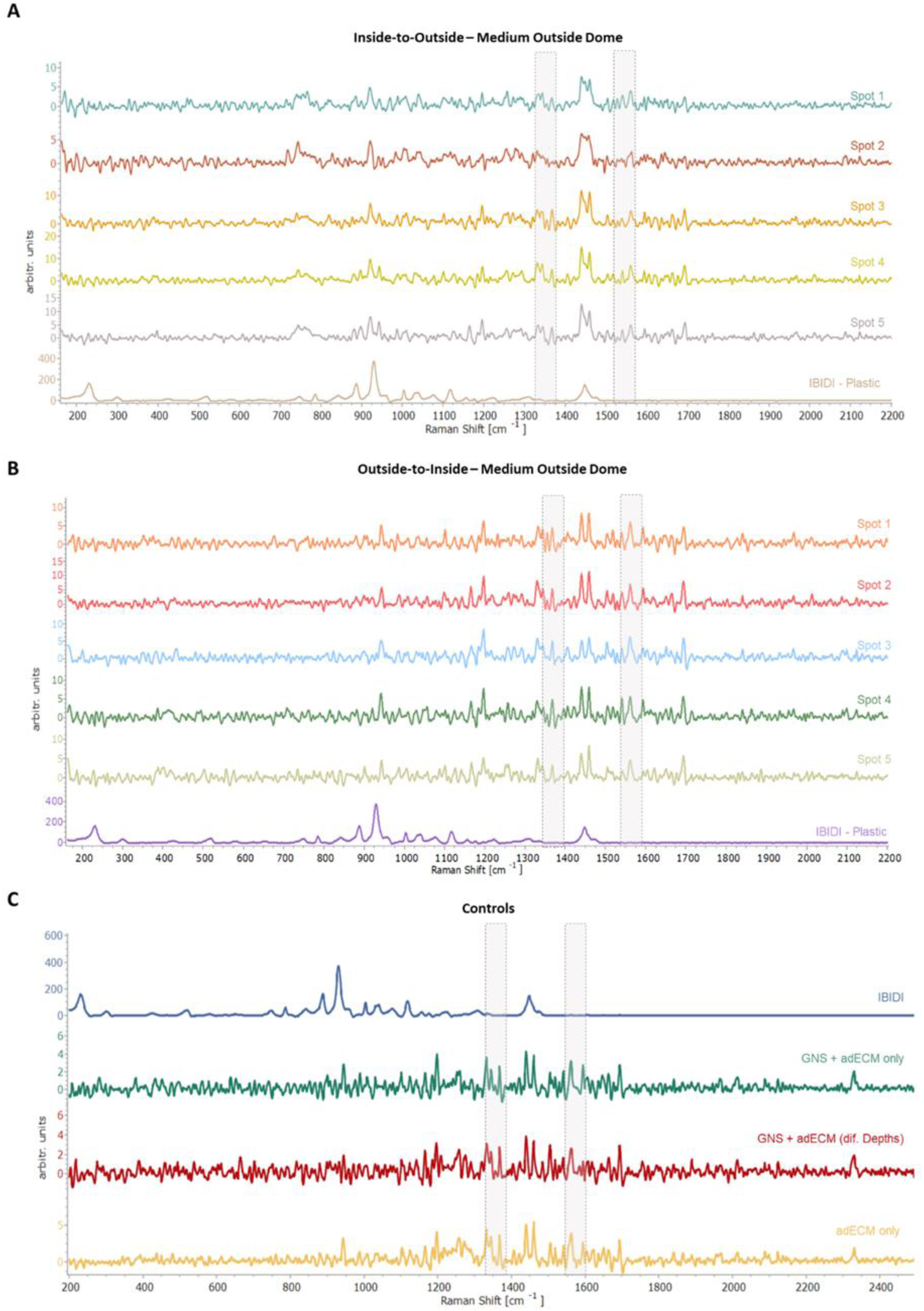
SERS spectra obtained from *in situ* SERS measurements of the medium outside the dome using the strategy: A) inside-to-outside; B) outside-to-inside and C) some controls composed by IBIDI plastic, GNS + adECM and adECM only. Five different spots were measured outside the dome. All spectra were acquired using an inverted confocal Raman microscope: 785 nm, laser power: 45mW and with a 50x objective, 2s and 10 acquisition.

### S11. Mapping of the different strategies used with 1-NAT in the domes

**Figure S11.**
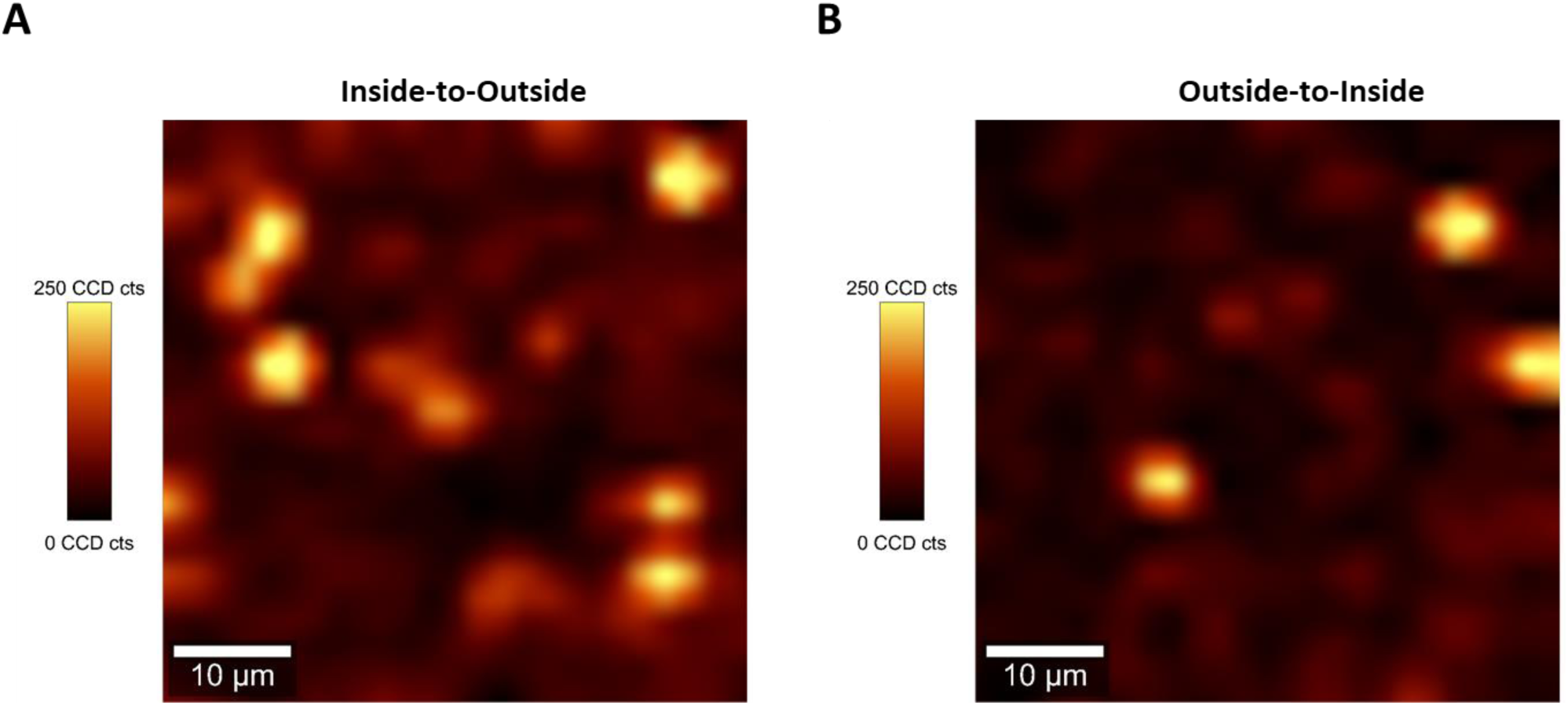
SERS mapping obtained from *in situ* SERS measurements in domes using the strategy: A) inside-to-outside and B) outside-to-inside. For both mapping the specific peak of 1-NAT 1,372 cm−1 was used demonstrating the distribution of the 1-NAT inside the dome. Both mapping were acquired using an inverted confocal Raman microscope. Mapping area: 50 µm x 50 µm, 2 µm step size, 0.5-second acquisition time, with the corresponding intensity colour scale. The specific peak from 1-NAT at 1,372 cm⁻¹ was selected for the maps, with a range of 35 cm⁻¹. Laser: 785 nm; Laser power: 45 mW; Objective: 50x.

## Notes

### Competing Interest Statement

The authors have declared no competing interest.

## References

1. Arnold M, Morgan E, Rumgay H, et al. Current and future burden of breast cancer: Global statistics for 2020 and 2040. The Breast. 2022;66:15–23. doi:10.1016/j.breast.2022.08.010

2. Liu Z, Chen J, Ren Y, et al. Multi-stage mechanisms of tumor metastasis and therapeutic strategies. Signal Transduct Target Ther. 2024;9(1):270. doi:10.1038/s41392-024-01955-5

3. Fares J, Fares MY, Khachfe HH, Salhab HA, Fares Y. Molecular principles of metastasis: a hallmark of cancer revisited. Signal Transduct Target Ther. 2020;5(1):28. doi:10.1038/s41392-020-0134-x

4. Liu M, Yang J, Xu B, Zhang X. Tumor metastasis: Mechanistic insights and therapeutic interventions. MedComm. 2021;2(4):587–617. doi:10.1002/mco2.100

5. Gilson P, Merlin JL, Harlé A. Deciphering Tumour Heterogeneity: From Tissue to Liquid Biopsy. Cancers (Basel). 2022;14(6):1384. doi:10.3390/cancers14061384

6. Dagogo-Jack I, Shaw AT. Tumour heterogeneity and resistance to cancer therapies. Nat Rev Clin Oncol. 2018;15(2):81–94. doi:10.1038/nrclinonc.2017.166

7. Turajlic S, Swanton C. Metastasis as an evolutionary process. Science (80-). 2016;352(6282):169–175. doi:10.1126/science.aaf2784

8. Paul ED, Huraiová B, Valková N, et al. The spatially informed mFISHseq assay resolves biomarker discordance and predicts treatment response in breast cancer. Nat Commun. 2025;16(1):226. doi:10.1038/s41467-024-55583-2

9. Marx V. Method of the Year: spatially resolved transcriptomics. Nat Methods. 2021;18(1):9–14. doi:10.1038/s41592-020-01033-y

10. Williams CG, Lee HJ, Asatsuma T, Vento-Tormo R, Haque A. An introduction to spatial transcriptomics for biomedical research. Genome Med. 2022;14(1):68. doi:10.1186/s13073-022-01075-1

11. Tivey A, Lee RJ, Clipson A, et al. Mining nucleic acid “omics” to boost liquid biopsy in cancer. Cell Reports Med. 2024;5(9):101736. doi:10.1016/j.xcrm.2024.101736

12. Lianidou E, Pantel K. Liquid biopsies. Genes, Chromosom Cancer. 2019;58(4):219–232. doi:10.1002/gcc.22695

13. Youk J, Kwon HW, Kim R, Ju YS. Dissecting single-cell genomes through the clonal organoid technique. Exp Mol Med. 2021;53(10):1503–1511. doi:10.1038/s12276-021-00680-1

14. Chang X, Zheng Y, Xu K. Single-Cell RNA Sequencing: Technological Progress and Biomedical Application in Cancer Research. Mol Biotechnol. 2024;66(7):1497–1519. doi:10.1007/s12033-023-00777-0

15. Zhang L, Lee M, Maslov AY, Montagna C, Vijg J, Dong X. Analyzing somatic mutations by single-cell whole-genome sequencing. Nat Protoc. 2024;19(2):487–516. doi:10.1038/s41596-023-00914-8

16. Habanjar O, Diab-Assaf M, Caldefie-Chezet F, Delort L. 3D Cell Culture Systems: Tumor Application, Advantages, and Disadvantages. Int J Mol Sci. 2021;22(22):12200. doi:10.3390/ijms222212200

17. Slater, K., Partridge, J. & Nandivada H. Tuning the Elastic Moduli of Corning Matrigel and Collagen I 3D Matrices by Varying the Protein Concentration. (Corning, 2019). https://www.corning.com/catalog/cls/documents/application-notes/CLS-AC-AN-449.pdf

18. Prince E, Cruickshank J, Ba-Alawi W, et al. Biomimetic hydrogel supports initiation and growth of patient-derived breast tumor organoids. Nat Commun. 2022;13(1):1466. doi:10.1038/s41467-022-28788-6

19. Schmidt DR, Patel R, Kirsch DG, Lewis CA, Vander Heiden MG, Locasale JW. Metabolomics in cancer research and emerging applications in clinical oncology. CA Cancer J Clin. 2021;71(4):333–358. doi:10.3322/caac.21670

20. Merteroglu M, Santoro MM. Exploiting the metabolic vulnerability of circulating tumour cells. Trends in Cancer. 2024;10(6):541–556. doi:10.1016/j.trecan.2024.03.004

21. Neuling NR, Allert RD, Bucher DB. Prospects of single-cell nuclear magnetic resonance spectroscopy with quantum sensors. Curr Opin Biotechnol. 2023;83:102975. doi:10.1016/j.copbio.2023.102975

22. Barshutina M, Arsenin A, Volkov V. SERS analysis of single cells and subcellular components: A review. Heliyon. 2024;10(18):e37396. doi:10.1016/j.heliyon.2024.e37396

23. Abalde-Cela S, Carregal-Romero S, Coelho JP, Guerrero-Martínez A. Recent progress on colloidal metal nanoparticles as signal enhancers in nanosensing. Adv Colloid Interface Sci. 2016;233:255–270. doi:10.1016/j.cis.2015.05.002

24. Xiao Y, Kim D, Dura B, et al. Ex vivo Dynamics of Human Glioblastoma Cells in a Microvasculature-on-a-Chip System Correlates with Tumor Heterogeneity and Subtypes. Adv Sci. 2019;6(8). doi:10.1002/advs.201801531

25. LeSavage BL, Suhar RA, Broguiere N, Lutolf MP, Heilshorn SC. Next-generation cancer organoids. Nat Mater. 2022;21(2):143–159. doi:10.1038/s41563-021-01057-5

26. Crapo PM, Gilbert TW, Badylak SF. An overview of tissue and whole organ decellularization processes. Biomaterials. 2011;32(12):3233–3243. doi:10.1016/j.biomaterials.2011.01.057

27. Neishabouri A, Soltani Khaboushan A, Daghigh F, Kajbafzadeh AM, Majidi Zolbin M. Decellularization in Tissue Engineering and Regenerative Medicine: Evaluation, Modification, and Application Methods. Front Bioeng Biotechnol. 2022;10. doi:10.3389/fbioe.2022.805299

28. Ohguro H, Watanabe M, Sato T, et al. Application of Single Cell Type-Derived Spheroids Generated by Using a Hanging Drop Culture Technique in Various In Vitro Disease Models: A Narrow Review. Cells. 2024;13(18):1549. doi:10.3390/cells13181549

29. Hanahan D, Weinberg RA. Hallmarks of cancer: the next generation. Cell. 2011;144(5):646–674. doi:10.1016/j.cell.2011.02.013

30. Hrelescu C, Sau TK, Rogach AL, et al. Selective Excitation of Individual Plasmonic Hotspots at the Tips of Single Gold Nanostars. Nano Lett. 2011;11(2):402–407. doi:10.1021/nl103007m

31. Däster S, Amatruda N, Calabrese D, et al. Induction of hypoxia and necrosis in multicellular tumor spheroids is associated with resistance to chemotherapy treatment. Oncotarget. 2017;8(1):1725–1736. doi:10.18632/oncotarget.13857

32. Nunes AS, Barros AS, Costa EC, Moreira AF, Correia IJ. 3D tumor spheroids as in vitro models to mimic in vivo human solid tumors resistance to therapeutic drugs. Biotechnol Bioeng. 2019;116(1):206–226. doi:10.1002/bit.26845

33. Zhuang P, Chiang YH, Fernanda MS, He M. Using Spheroids as Building Blocks Towards 3D Bioprinting of Tumor Microenvironment. Int J Bioprinting. 2024;7(4):444. doi:10.18063/ijb.v7i4.444

34. Sun X, Kaufman PD. Ki-67: more than a proliferation marker. Chromosoma. 2018;127(2):175–186. doi:10.1007/s00412-018-0659-8

35. Laurent J, Frongia C, Cazales M, Mondesert O, Ducommun B, Lobjois V. Multicellular tumor spheroid models to explore cell cycle checkpoints in 3D. BMC Cancer. 2013;13:73. doi:10.1186/1471-2407-13-73

36. Yakavets I, Francois A, Benoit A, Merlin JL, Bezdetnaya L, Vogin G. Advanced co-culture 3D breast cancer model for investigation of fibrosis induced by external stimuli: optimization study. Sci Rep. 2020;10(1):21273. doi:10.1038/s41598-020-78087-7

37. Di Rosa F, Cossarizza A, Hayday AC. To Ki or Not to Ki: Re-Evaluating the Use and Potentials of Ki-67 for T Cell Analysis. Front Immunol. 2021;12. doi:10.3389/fimmu.2021.653974

38. Sutherland RL, Hall RE, Taylor IW. Cell proliferation kinetics of MCF-7 human mammary carcinoma cells in culture and effects of tamoxifen on exponentially growing and plateau-phase cells. Cancer Res. 1983;43(9):3998–4006. http://www.ncbi.nlm.nih.gov/pubmed/6871841

39. Prado-Garcia H, Campa-Higareda A, Romero-Garcia S. Lactic Acidosis in the Presence of Glucose Diminishes Warburg Effect in Lung Adenocarcinoma Cells. Front Oncol. 2020;10. doi:10.3389/fonc.2020.00807

40. Contorno S, Darienzo RE, Tannenbaum R. Evaluation of aromatic amino acids as potential biomarkers in breast cancer by Raman spectroscopy analysis. Sci Rep. 2021;11(1):1698. doi:10.1038/s41598-021-81296-3

41. Becerril-Castro IB, Calderon I, Pazos-Perez N, et al. Gold Nanostars: Synthesis, Optical and SERS Analytical Properties. Anal Sens. 2022;2(3). doi:10.1002/anse.202200022

42. Turkevich J, Stevenson PC, Hillier J. A study of the nucleation and growth processes in the synthesis of colloidal gold. Discuss Faraday Soc. 1951;11:55. doi:10.1039/df9511100055

